# Thermoresponsive, Shear-Thinning Supramolecular Hydrogels from Peptide–Polymer Conjugates for Bioinspired 3D Cell Culture

**DOI:** 10.64898/2026.09.04.749227

**Authors:** Breanna M. Huntington, Johannes A. Dresel, Maren Schweitzer, Eric M. Furst, Pol Besenius, April M. Kloxin

## Abstract

Physically assembled, supramolecular hydrogels often are formed by weak bonds, producing useful viscoelastic properties that are reminiscent of soft tissues within the human body yet presenting challenges in stability under physiological conditions. In this work, we establish an approach for creating self-assembling, supramolecular hydrogels based on a peptide-polymer conjugate (PPC) as a three-dimensional (3D) cell culture platform. The viscoelastic properties of these hydrogels, including thermoresponsiveness and shear-thinning, were measured using rheometry. Building on these observations, we then developed a method for the encapsulation and culture of human wound healing cells, fibroblasts, within the PPC hydrogels, observing both excellent cell viability and material stability over time. The modulation of stiffness by temperature variation and the reversible nature of these hydrogels imparts unique properties for both cell encapsulation and harvesting and offers potential for a variety of biomedical applications, from 3D cell culture systems to therapeutic delivery.

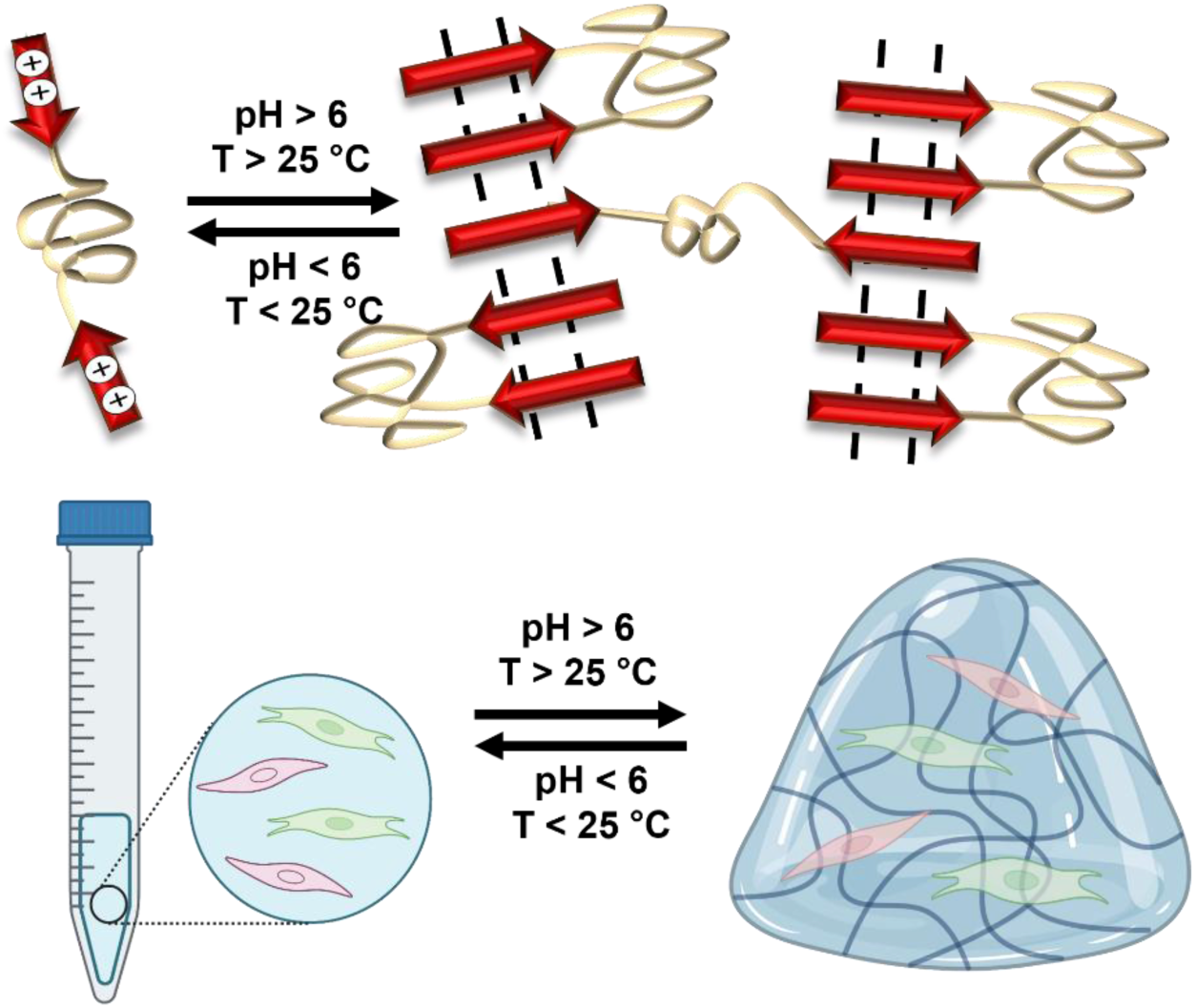

## Introduction

Physically crosslinked hydrogels have garnered significant attention in recent years due to their tunable viscoelastic properties, including shear thinning and self-healing in response to stimuli relevant for a range of applications.^1^ Their underlying molecular structure is crosslinked with non-covalent interactions between hydrophilic macromolecular building blocks, such as hydrogen bonding and π-π stacking.^1, 2^ Moreover, their ability to form non-covalent, reversible bonds often results in responsive viscoelastic properties. These tunable interactions not only impart desirable viscoelastic properties, but also enable controlled gelation and dissolution or degradation, allowing for subsequent processing and dynamic behavior in response to external stimuli such as changes in pH, ion concentration, temperature, or applied strain.^3^ One field of research with growing interest in those properties is three-dimensional (3D) cell encapsulation and culture systems and more broadly bioprinting, where biocompatible, thermoresponsive polymers can be processed through temperature-controlled transitions to create tissue-like 3D environments.^4, 5^ For example, thermoresponsive building blocks, such as those based on Pluronic F-127 and PNIPAM, have been explored; however, in these applications, nonphysiological temperature handling associated with thermally-induced gelation or phase separation can alter the water content, volume, and mechanical properties of the resulting hydrogel, which may complicate cell encapsulation and culture.^6–9^ Further, the mechanical properties of unmodified systems can be difficult to tune independent of polymer concentration and transition temperature, limiting their ability to recapitulate specific soft tissue microenvironments. More broadly, the mechanical strength and stability of traditional physically-assembled constructs often are limited under physiologically relevant conditions, presenting challenges in maintaining hydrogel structure and shape over time in biological applications of interest.^3,6^ These limitations highlight the need for physically crosslinked hydrogel systems that combine dynamic responsiveness with improved structural robustness for their use and interfacing with biological systems.

The reversible interactions that underlie physically crosslinked hydrogels often also support the formation of designed hierarchical structures, including secondary and tertiary formations like α-helices, β-sheets, coiled-coils, and fibers.^5, 10–12^ These structures hold considerable promise in biomedical applications, with opportunities to serve as responsive tissue mimics upon balancing their stability and viscoelastic properties under physiologically relevant conditions.^13–21^ To realize the potential of specific physically crosslinked systems in applications of interest, approaches are needed for establishing relevant material formulations and translating them to address outstanding needs, including for 3D cell culture and therapeutic delivery.^3, 11, 22, 23^ For example, drug-loaded physically-crosslinked injectable hydrogels have been shown to increase the residence time of the drug as well as the overall therapeutic outcome compared to conventional administration, where their pH-responsive behavior can be exploited for triggered, local release at tumor sites.^24–26^ Building blocks for the creation of well-defined, responsive supramolecular hydrogels with tailorable properties remain a prominent need for deployment of such materials in a range of cellular microenvironments.

Peptides are the fundamental building blocks of proteins and consist of amino acids linked through amide bonds whose side chains impart physical interactions. These guide their assembly into secondary and higher ordered structures, which can be exploited for the formation of peptide-based hydrogels with designed structures and properties.^27^ However, the use of naturally-occurring peptides for the creation of such hierarchical structures can be limited by the restricted tunability of key properties, such as solubility, mechanical strength, and responsiveness to external stimuli.^28^ Chemical modifications of specific amino acids, as well as the *de novo* design of new engineered peptides, can be employed for the development of hydrogel-based materials with properties inspired by those of naturally-occurring peptides.^29, 30^ Aside from imparting biocompatibility and biodegradability, the peptide structure of the hydrogel can offer precise control of biochemical and biophysical properties through the underlying sequence of amino acids. Scalable strategies that leverage the programmability of peptide-based assemblies while overcoming the limitations of naturally occurring systems remain an important objective for hydrogel design.

The approach used in this work combines the scalability and ease of access of synthetic polymers with the tunability and unique characteristics of peptides using peptide-polymer conjugate (PPC) building blocks.^31^ PPCs consist of peptide sequences coupled with synthetic polymers, thus mitigating the potential insolubility or instability of specific assembling peptides while harnessing their responsive, molecularly-engineered hierarchical structures.^32, 33^ Here, we selected a PPC comprising of a difunctionalized polyethylene glycol (PEG) chain flanked by two heptapeptides containing repeating units of histidine and phenylalanine residues.^34^ Originally developed by Otter *et al.*, the physical properties of these PPC hydrogels have been probed using both experimental and computational methods for an understanding of how the PEG chain improves flexibility of the PPC in aqueous environments and the peptide segments impart physical assembly via hydrogen bonding and π-π stacking.^34, 35^ The properties imparted by the telechelic ABA peptide-polymer-peptide design make them attractive for 3D cell culture, with the potential for introducing thermal and strain responsiveness to the synthetic matrix produced with these water-soluble assembling building blocks.

Specifically, in this work, we examined the properties of these telechelic ABA peptide-polymer-peptide hydrogels under physiologically relevant conditions and their use as a platform for 3D cell culture. We first conducted a series of shear rheometry measurements to evaluate the mechanical, viscoelastic, and temperature responsive properties of the hydrogels. These PPCs were observed to form weak hydrogels at room temperature and exhibit increased moduli and good stability at physiological temperatures, as well as reversible thermoresponsiveness for rapid dissolution upon cooling, offering unique advantages for 3D cell culture and harvesting. Based on these observations, we hypothesized that cells could be readily encapsulated in these PPC hydrogels for stable 3D cell cultures with lung tissue inspired moduli leveraging the reagent-free, thermoreversible assembly mechanism for both encapsulation with good viability, phenotypic maintenance of cell types that are sensitive to their mechanical microenvironment (e.g., human fibroblasts), and efficient, non-destructive cell recovery. To test this, a workflow was established for encapsulating wound healing cells (human fibroblasts) within the hydrogels with relevant stability for 3D cell culture. Fibroblasts were used given their prevalence in tissues throughout the body and their importance in healthy and maladaptive wound healing processes.^36, 37^ We then monitored the viability of the encapsulated fibroblasts to assess relevance for cell culture over time, both *in-situ* and upon thermally responsive harvesting of the culture cells. Overall, we found that the PPC hydrogels had useful properties for cell encapsulation and culture and, under the processing conditions and workflows established here, promoted good cellular viability and function over time. These findings establish the designed thermoresponsive PPC hydrogels as an adaptable biomaterial platform for 3D cell culture and provide a foundation for future studies investigating their use with a range of cell types for dynamic tissue models and for therapeutic delivery applications.

## Results and Discussion

To probe applicability of the PPC hydrogels for 3D bioinspired cell culture, we analyzed the mechanical and viscoelastic properties of the assembled hydrogels and developed a method to three-dimensionally encapsulate a human pulmonary fibroblast cell line (CCL 151) within the gels (**Figure 1**). Block copolymers are known to have unique viscoelasticity that can provide tunable properties to synthetic ECMs for bioinspired cell cultures.^38–40^ Additionally, these polymers often exhibit unique thermoresponsiveness, owing to changes in hydrophobicity and hydrophilicity with varying temperature, an advantageous feature that can be exploited in many biomedical applications.^41, 42^ Here, we deployed an ABA-type PPC comprised of a telechelic PEG chain with heptapeptide end groups rich in histidine and phenylalanine. This PPC was selected for its aqueoussolubility and ability to undergo reversible, pH-dependent self-assembly via hydrogen bonding and π–π stacking in aqueous conditions, which we hypothesized would be relevant for creating thermally and mechanically responsive synthetic matrices for 3D cell culture.^34, 35^ This PPC was synthesized by established protocols and characterized with NMR, MALDI, and GPC to verify the identity and purity of the building block, as detailed in the Supporting Information (**Figures S1 and S2**).^34, 35^

**Figure 1.**
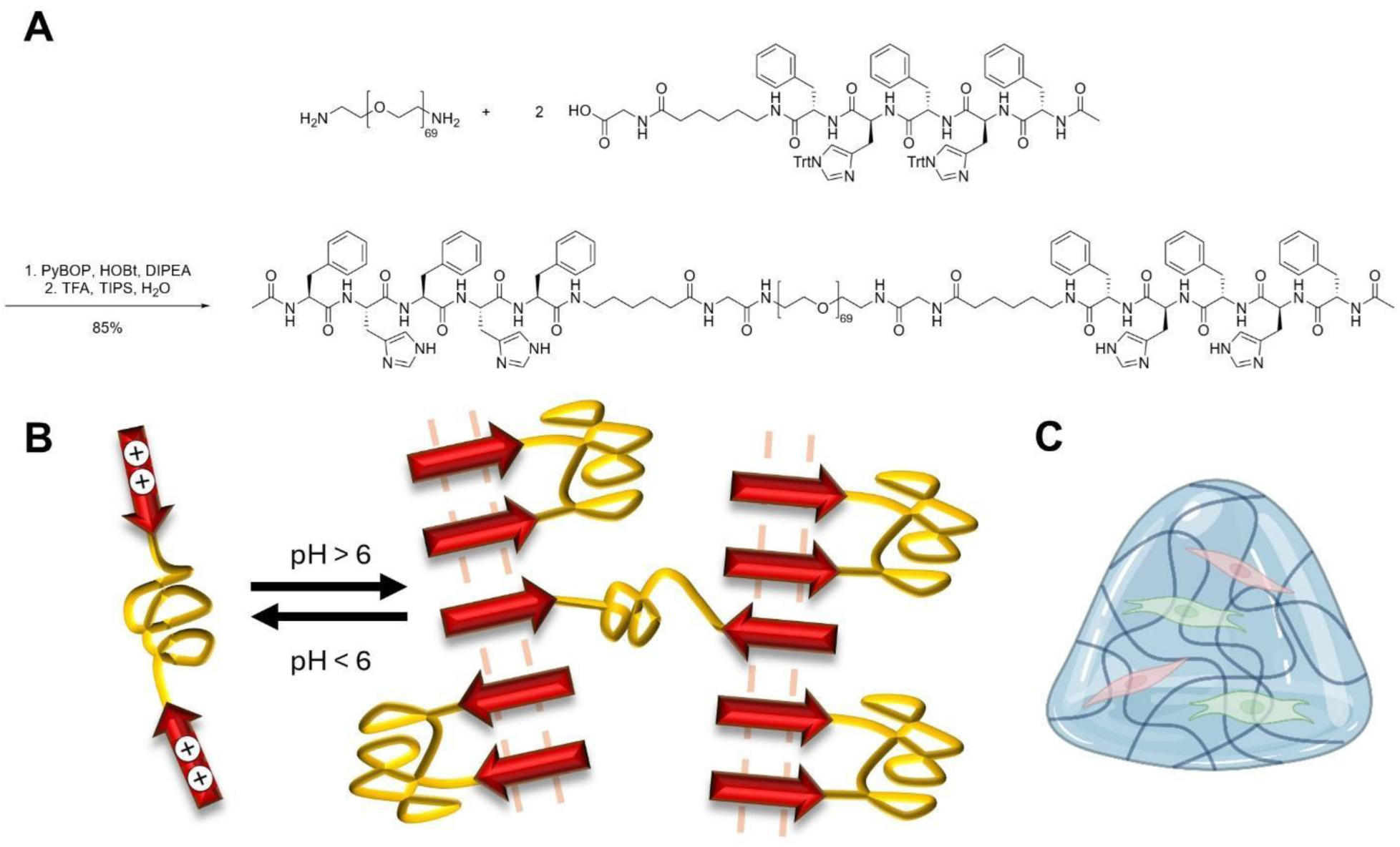
Overview of approach. (A) Synthesis of the polymer-peptide conjugate (PPC). The peptide (Phe-His-Phe-His-Phe-Aminohexanoic acid-Gly-OH, [Ac-FHFHF-Ahx-G-OH]) was synthesized by solid phase peptide synthesis and then coupled to PEG *bis*-amine (M_n_ ∼ 3000 g/mol) to form the PPC. (B) Schematic of pH-induced self-assembly for hydrogel formation. (C) Schematic representation of the 3D cell culture model.

We used rheometry to assess the viscosity, shear thinning behavior, and responsiveness to changes in temperature.^43^ First, to examine the shear thinning behavior and thereby the potential for processing of the PPC hydrogels (e.g., for cell encapsulation, injectability), we subjected them to a flow curve study (**Figures 2A and S3**). Protocols were established for forming hydrogels under physiologically relevant conditions and *in situ* on the rheometer for these studies. Briefly, 1% wt/vol PPC hydrogel precursor solution was prepared by dissolving the PPC in PBS at a pH of 2.4. The pH of the PPC was then brought up to 7.0 using 0.1 M NaOH. The hydrogel precursor solution was pipetted on to the rheometer and allowed to form *in-situ* for 30 minutes. A viscosity flow study was conducted at a shear rate from 0.01/s to 30.0/s, and we observed that, as the shear rate increased, the viscosity of the gel decreased, indicating that the gels were shear thinning.^44^

**Figure 2.**
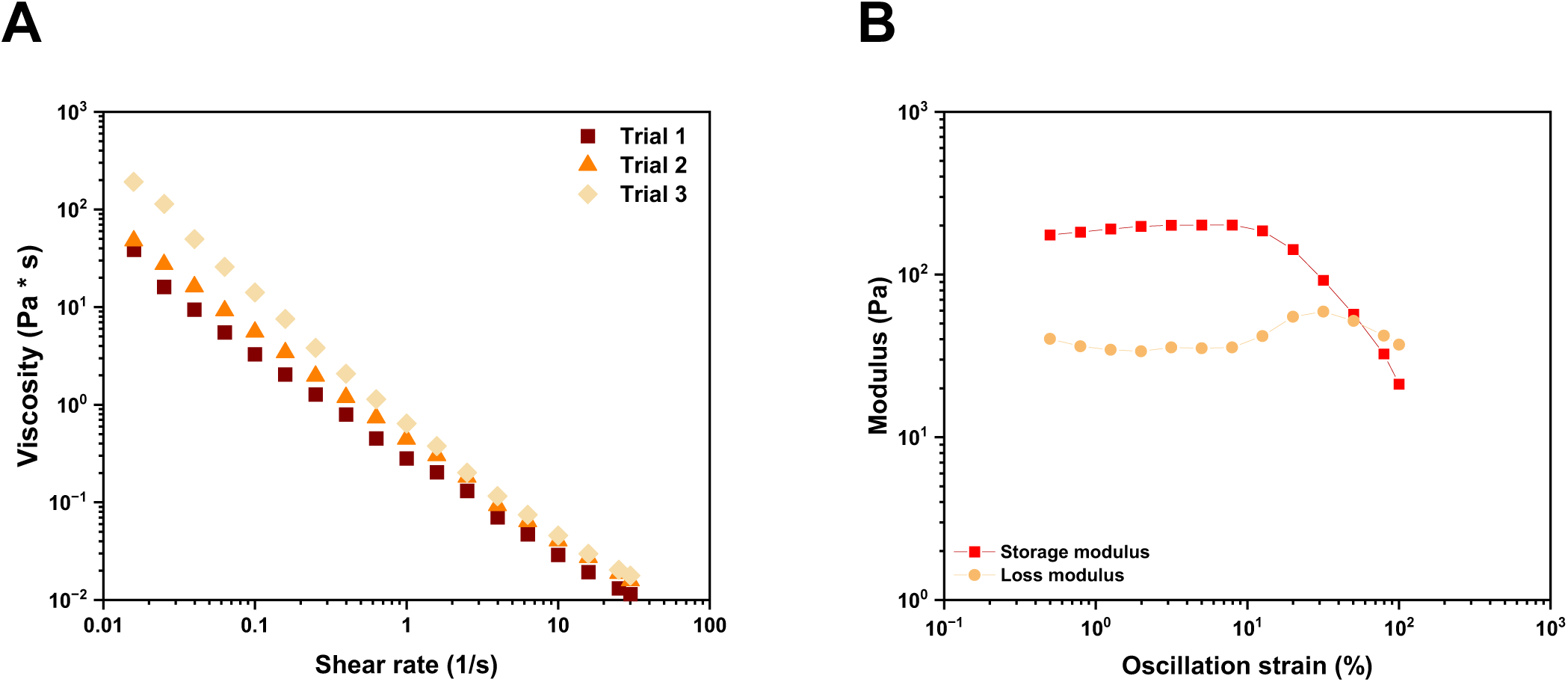
Characterization of non-linear elastic, shear-thinning properties using rheometry. (A) Viscosity versus shear rate for hydrogels. (B) Strain amplitude sweep of hydrogels.

To further study the extent of viscoelastic behavior, we formed hydrogels *in-situ* on the rheometer and subjected them to an amplitude sweep from 0.1% to 100% strain (**Figures 2B and S5**). We observed crossover of the storage modulus (G’) and loss modulus (G”), indicating a transition from gel to sol (gel-like to liquid-like properties) at high strains, further supporting viscoelastic behavior for these physically crosslinked hydrogels. This viscoelastic behavior is beneficial for material processing including in biological applications (e.g., gel-based encapsulation, injectable delivery vehicles).^39, 45^ Notably, the storage modulus observed within the linear viscoelastic regime under low strain conditions is ∼ 0.2 kPa, a similar order of magnitude to the lower moduli reported for the lung parenchyma amongst other soft tissues.^46^ Further, the oscillatory strain at which thinning is observed is similar to that reported for lung tissues.^46^ Overall, a PPC composition was identified with properties relevant to and inspired by soft tissues like those found in the lung and that enabled processing by leveraging viscoelastic properties.

The physical crosslinking mechanism of these PPC hydrogels was hypothesized to impart not only viscoelasticity for processing, but also thermal responsiveness in ranges of potential relevance for biological applications. To assess responsiveness to temperature, the hydrogels were subjected to a series of temperature studies, examining modulation of mechanical properties with both increases and decreases in temperature within ranges compatible with cells and tissues (e.g., 4 °C to 37 °C) (**Figures 3, S6, and S7**).

**Figure 3.**
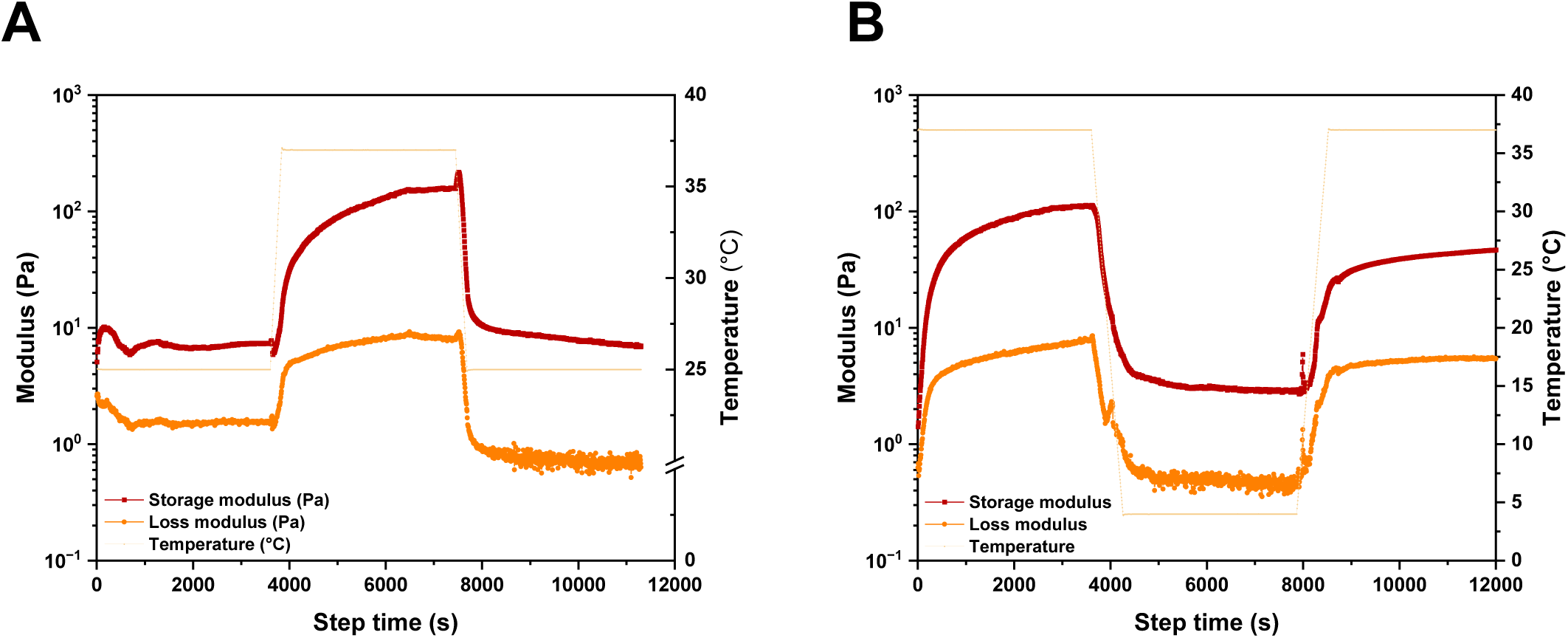
Rheometric assessment of temperature responsiveness. (A) Hydrogel properties upon heating and cooling cycles: the gels were allowed to form at 25 °C, temperature was ramped up to 37 °C, held steady for 1 hr, and ramped down again to 25 °C. (B) Hydrogel properties upon cooling and heating cycles: gels were allowed to form at 37 °C, temperature was ramped down to 4 °C, held steady for 1 hr, and ramped up again to 37 °C.

Hydrogels were prepared as before at 1% wt/vol and allowed to form *in-situ* on the rheometer, using a cone and plate geometry with a solvent trap to prevent sample drying over the experimental time course. To first study how the hydrogel responds to increases in temperature (**Figure 3A**), hydrogels were formed at room temperature (1 hr at 25 °C) and then ramped to physiological temperature (1 hr at 37 °C). Upon increasing temperature to 37 °C, the moduli (storage G’ and loss G”) of the gel increased while retaining elastic properties (G’ > G”), supporting the potential of these hydrogels for 3D cell culture and other biological applications with gel handling at room temperature followed by use at physiological temperature. We speculate that the increase in storage and loss moduli upon heating from 25 to 37 °C reflects temperature-dependent stabilization of the supramolecular network: for example, heating is known to enhance hydrophobic and aromatic association among the phenylalanine residues and may thereby increase the number or lifetime of transient junctions between the telechelic PEG chains.^47–50^ Because the network is maintained by noncovalent associations, these changes should be reversible upon cooling. Indeed, upon cooling to 25 °C, the modulus of the gel was observed to return to similar values as before the temperature ramp, supporting reversible thermoresponsiveness and the relevance of temperature cycling during processing of these materials in cytocompatible temperature ranges.

The hydrogels were subsequently subjected to decreases in temperature (**Figure 3B**) where we hypothesized that cooling would further decrease their storage and loss moduli by shifting the balance of noncovalent interactions within the network (e.g., increasing hydration and weakening hydrophobic association of the peptide end groups),^47, 51^ with the potential for a transition to a low-viscosity sol at 4 °C. In particular, dissolution of the gel at lower temperatures could impart beneficial properties for this system for use in 3D cell culture applications by allowing for easier experimental workflows that do not require the addition of external reagents for cell harvesting. Hydrogels were again formed *in-situ* at physiologically relevant conditions (1 hr at 37 °C) and then the temperature was ramped down (1 hr at 4 °C). Upon cooling, a significant decrease in modulus was observed. Further, through gentle shaking of cooled samples, the hydrogel was completely dissolved, transitioning from gel to sol. This dissolution upon cooling to a standard refrigeration temperature used in cell processing workflows is a rather unique and beneficial property of these supramolecular hydrogels for 3D cell culture. Such property modulation upon cooling offers the potential for a rapid and facile method to easily harvest cells from the hydrogels, for use either in cellular analyses or re-encapsulation in a new hydrogel without using added passaging reagents.^41^ The temperature was then ramped back up to 37 °C and similar hydrogel moduli to the initial condition were observed. We speculate that the slightly lower modulus relative to initial upon reheating is due to changes in the transient junctions between the telechelic PEG chains^47–50^ within the supramolecular network that occur with thermal processing. Overall, the rheometric assessment of the PPC hydrogels and their temperature responsiveness demonstrated that formation and dissolution take place in a physiologically relevant temperature window and moduli can be thermally modulated.

Next, we investigated whether the formation mechanism of these hydrogels would be compatible with encapsulation of wound healing cells (human lung fibroblasts (CCL 151s)) of relevance in 3D cell culture and delivery. Prior to encapsulating cells, stability of the hydrogels in different cell culture media was qualitatively probed. Hydrogels were formed in wells of a chamber slide at 37 °C, and standard growth medium (Kaighn’s F12K with 10% fetal bovine serum (FBS)) was then added at 37 °C. The FBS-containing growth medium led to rapid dissolution of the hydrogels. We hypothesized that components of serum may interfere with the hydrogen bonding mechanism of the physical crosslinks underlying these gels. Hydrogel stability in a commercially available, serum-free complete growth media (fibroblast basal medium (ATCC, PCS-201-030) with added serum-free fibroblast growth kit (ATCC, PCS-201-040)) was then evaluated. Importantly, hydrogels maintained their shape upon the application of this serum-free complete growth media and did not dissolve over 5 days of incubation at 37 °C.

For cell encapsulation and 3D culture, PPC-hydrogel precursor solution was prepared as before, dissolving at 1% wt/vol in PBS at pH 2.4. Cells were harvested from tissue culture flasks using trypsin and centrifuged to form a pellet, and the media was decanted. The pH of the hydrogel precursor was brought up to 7.0 using 0.1 M NaOH, and the cell pellet was resuspended in the precursor solution. Hydrogel precursor solution with suspended cells was pipetted into wells of chamber slides and incubated at 37 °C for 30 min before fresh serum-free complete media warmed to 37 °C was added on top. To examine cellular viability in this system, we used a membrane integrity assay (live/dead cytotoxicity assay) at time points of interest over a 5-day time frame and imaged with confocal microscopy (**Figures 4A, 4B and S8**). We found that approximately 90% of cells encapsulated in these hydrogels remain viable over the 5-day time-period, supporting the relevance of these hydrogels for 3D cell culture applications. Note, while our dynamic PPC system was permissive to cell culture and relevant assays (e.g., live/dead cytotoxicity assay), some reagents including nuclear stains and proteins used in immunostaining and biochemical assays (e.g. Hoechst, Alamar Blue, bovine serum albumin (BSA)) led to hydrogel dissolution. While certain biological reagents were observed to trigger hydrogel dissolution, this responsiveness can be viewed as a programmable design feature: for example, as these molecules act as chemical triggers, they may enable staged cell harvesting or the controlled release of encapsulated payloads, offering a degree of responsiveness to small molecules and non-enzymatic proteins not typically found in synthetic matrices. Beyond basic viability, maintaining and monitoring cell phenotype in a 3D microenvironment is critical. We demonstrated the utility of this platform for non-invasive phenotypic monitoring by encapsulating a lentiviral-based fluorescent reporter fibroblast line (constitutively expressing red, conditionally expressing green upon alpha smooth muscle actin upregulation) (Figure S10). This approach allowed for the real-time imaging of alpha-smooth muscle actin (α-SMA) expression, a key marker of myofibroblast activation, by fibroblasts in these 3D cultures without the need for destructive end-point immunostaining or the addition of external reagents (**Figure S10**).^52, 53^

**Figure 4.**
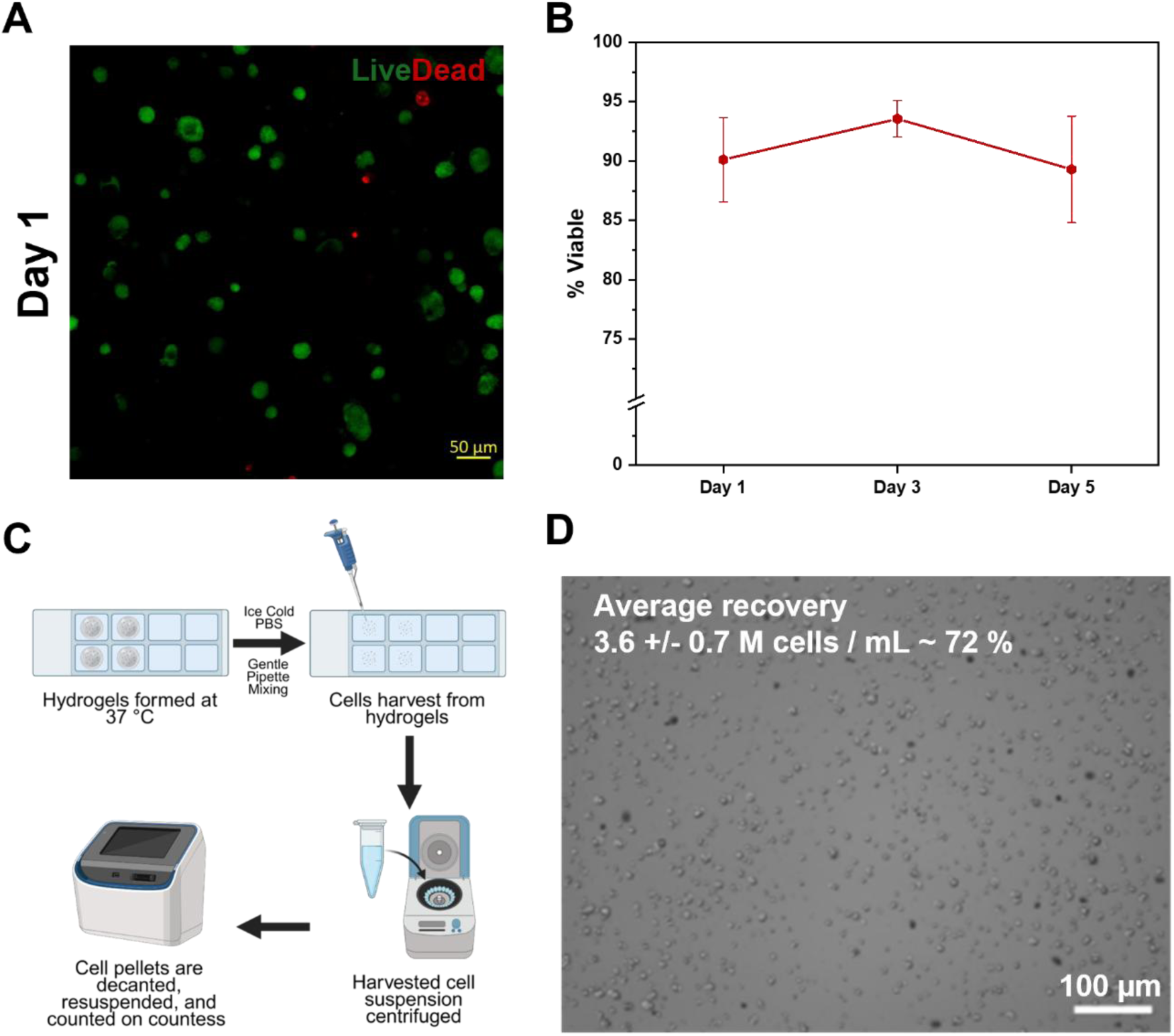
3D cell culture and cell harvesting. (A) Viability of cells encapsulated in 3D in the hydrogels was assessed using a Live/Dead assay. Representative projection of confocal Z-stacks (thickness ∼ 100 µm) of live (green) / dead (red) cells at day 1 after encapsulation. (B) Quantification of cellular viability over a 5 day time period in 3D culture through image analysis using Volocity. (C) Process for cell harvesting exploiting thermoresponsive properties. (D) Representative image of recovered cells that were harvested from 3D culture and quantified with a Countess cell counter. Over 4 replicates (each containing 4 harvested samples), an average of 3.6 M cells/mL +/- 0.7 M cells/mL was collected, resulting in ∼ 72% of cells recovered based on the initial seeding density of cells encapsulated in the hydrogels.

Last, we leveraged the reversible thermoresponsive nature of the PPC hydrogel network, as described in the shear rheology characterization paragraph, to establish a straightforward protocol for cell harvesting (**Figure 4C, 4D and S9**). To this end, 3D culture samples were removed from the incubator, media was removed, and a small volume (∼ 100 µL) of ice-cold PBS was added to the well. This protocol resulted in immediate dissolution of the hydrogel, followed by recovery of the cells from the sol by centrifugation. To make one technical replicate, 4 samples were pooled together to provide a visible cell pellet and ensure a sufficient number of cells for analysis using a Countess cell counter. By applying ice-cold PBS to trigger dissolution, we recovered cells at a mean density of 3.6 M cells/mL, representing a ∼72% recovery based on the initial seeding density. Importantly, this temperature-driven harvesting protocol avoids the use of proteolytic enzymes like trypsin, which are known to degrade cell surface proteins, thereby preserving the cellular state for sensitive downstream applications (e.g., flow cytometry or transcriptomics). These properties will open future avenues in the design of dynamic ECM mimics and bioinspired cell culture materials that facilitate cell passaging, harvesting, or injection which is not possible to the same extent with conventional covalently crosslinked 3D matrices. Overall, a significant advantage of this thermoresponsive system is the ability to achieve reagent-free cell harvesting with high efficiency.

## Conclusion

In this work, we have demonstrated that bulk rheometric techniques can be successfully used to characterize various viscoelastic properties of PPC hydrogels, including their viscosity in response to increasing shear rate, their shear thinning behavior in response to applied strain, and their reversible temperature responsiveness, establishing a framework for evaluating such materials for use in biological applications and highlighting temperature ranges for deploying these specific PPC materials in 3D culture. Based on the thermoresponsive viscoelastic properties of the PPCs, we developed a method to encapsulate and culture human fibroblasts and observed good cellular viability over a 5-day timespan in these 3D cultures, achieving good stability and lung tissue inspired moduli at 37 °C and facile cell harvesting at 4 °C. The combination of shear-thinning behavior and reversible thermoresponsiveness positions these PPC hydrogels as promising candidates for advanced bioprinting applications: for example, where the material can protect cells during the high-shear extrusion process and rapidly stabilize upon contact with a heated stage.

Further, the increased moduli and stabilization at physiological temperatures make this system attractive for use as an injectable cell delivery vehicle, with the potential to provide a protective, bioinspired matrix that supports cellular phenotype and function. Our findings suggest that the PPC hydrogels offer an innovative option for 3D bioinspired cell cultures and could open avenues for further applications in 3D cell culture systems for cell expansion and delivery (e.g., that allow for facile cell passaging, harvesting, or injection in these 3D matrices). Building from these studies, future work could include new engineered PPCs that physically or covalently integrate bioactive groups (e.g., cell-binding peptides) to modulate cellular activities in 3D cultures. Additionally, tuning the molecular structure of the PPC opens opportunities to program the mechanical properties of the resulting hydrogel. Further, the thermal and shear responsiveness of the PPC synthetic matrix may prove useful in future biomedical applications such as injectable therapeutic delivery vehicles.

## Associated Content

### Supplementary information

The supplementary information (SI) is available free of charge. The SI includes synthesis and characterization data of the PPC (detailed synthetic procedure, NMR, MALDI-ToF, rheology) as well as additional imaging and data for cell experiments.

### Funding

This work was supported by grants for related work from a National Institutes of Health (NIH) Director’s New Innovator Award with grant number DP2HL152424 (Kloxin) and the National Science Foundation (NSF) through the University of Delaware Materials Research Science and Engineering Center (DMR-2011824). Funding from the DFG (Deutsche Forschungsgemeinschaft) is also acknowledged: P.B. is PI and M.S. an associate student of the GRK 2516 (Project No. 405552959); P.B. is a PI of the SFB 1552 (Project No. 465145163). P.B. also acknowledges the European Research Council (ERC) under the European Union’s Horizon 2020 research and innovation program (ERC CoG SUPRAVACC 819856) for financial support. M.S. was the recipient of a doctoral fellowship from the Max Planck Graduate Center with the Johannes Gutenberg University Mainz (MPGC). B. H. was the recipient of the Schipper Fellowship from the Department of Chemical and Biomolecular Engineering at the University of Delaware and the Graduate Assistance in Areas of National Need (GAANN) fellowship funded through the U.S. Department of Education. Additionally, the authors acknowledge the use of facilities and instrumentation supported by the NSF through the University of Delaware Materials Research Science and Engineering Center (DMR-2011824) and the NIH National Institute of General Medical Sciences (NIGMS) through the Delaware COBRE (P20GM104316). BioRender.com was used for schematic creation within figures.

