## supplemental information for "Thermoresponsive, Shear-Thinning Supramolecular Hydrogels from Peptide–Polymer Conjugates for Bioinspired 3D Cell Culture"

**Materials and Methods**

***Part I: Synthesis of Polymer-Peptide Conjugate***

Materials: All solvents and reagents were obtained from commercial sources at highest grade and used without further purification. DMF and piperidine used during peptide synthesis were used in peptide grade quality. H_2_N-PEO(3000)-NH_2_ was purchased from *Rapp Polymere* (Tübingen, Germany). Acetic anhydride, TIPS, peptide-grade DMF and matrices for MALDI-ToF were purchased from *Sigma Aldrich* (St. Louis, MO, USA). Fmoc-protected amino acids, HBTU and PyBOP were purchased from *Carbolution* (St. Ingbert, Germany), aminohexanoic acid from *Acros Organics* (Geel, Belgium). DIPEA, DCM and toluene were purchased from *Thermo Fisher Scientific* (Waltham, MA, USA). HOBt was obtained from *abcr* (Karlsruhe, Germany), TFE from *FluoroChem* (Glossop, UK) and TFA from *TCI* (Tokyo, Japan). 2-Chlorotrityl resin and peptide-grade piperidine were purchased from *Iris Biotech* (Marktredwitz, Germany), DMSO-*d*_6_ from *Deutero* (Kastellaun, Germany) and diethyl ether from *VWR* (Radnor, PA, USA). Solid phase peptide synthesis was performed on a *CS136XT Peptide Synthesizer* from *CSBio Co.* (Menlo Park, CA, USA) using standard Fmoc-coupling protocols. ^1^H NMR spectroscopy was performed on an *Avance II 400* spectrometer (400 MHz), MALDI-ToF spectroscopy on an *autoflex©* MALDI-ToF spectrometer (concentration of 1 mg/mL), both from *Bruker* (Rheinstätten, Germany). Centrifugation was performed on a *VWR Mega Star 600* by *Avantor* (Darmstadt, Germany) in 50 mL *Falcon* tubes. Gel permeation chromatography was performed on an *Agilent 1100 Series* by *Agilent Technologies* (Santa Clara, CA, USA) in HFIP containing 3 g/L KTFA at 40 °C using PSS PFG 7 µM columns (100/1000 Å porosity) from *PSS Polymer Standards Services* (Mainz, Germany) a UV and RI detector. Poly(ethylene glycol) standards from *PSS Polymer Standards Services* (Mainz, Germany) were used for calibration.

Synthesis of the polymer peptide conjugate Ac-FHFHF(Ahx)G-PEO(3000)-G(Ahx)FHFHF-Ac:

(1) Preloading of the 2-chlorotrityl resin

Peptide synthesis was performed on 2-chlorotrityl resin. The resin was preloaded with the first amino acid: Fmoc-Gly-OH (761 mg, 2.56 mmol, 2.0 eq) was dissolved in DCM (10 mL) and added to 2-chlorotrityl resin (loading capacity 1.60 mmol/g, 800 mg, 1.28 mmol, 1.0 eq) in a Merrifield reactor. After DIPEA (446 μL, 2.56 mmol, 2.0 eq) was added and the reaction mixture agitated at room temperature (r.t.) for 5 min, another portion of DIPEA (446 μL, 2.56 mmol, 2.0 eq) was added and the reaction mixture agitated for 1 h. To inactivate non-functionalized sites on the resin, methanol (2 mL) was added, and the reaction mixture was agitated for 15 min at r.t. The Merrifield reactor was drained, and the resin consecutively washed with DCM (3 x), DMF (3 x) and DCM (3 x). The resin was used for peptide synthesis without further drying steps.

(2) Peptide synthesis

Peptide synthesis was performed on an automated solid phase peptide synthesizer. First, the preloaded resin was swollen in DCM for 1 h.

(2a) Fmoc-deprotection: The resin was covered with a solution of 20% piperidine in DMF and shaken for 10 min. After draining the cleavage solution, the step was repeated for 20 min. The resin was washed with DCM (2 x) and DMF (5 x).

(2b) Coupling (1.28 mmol scale): Amino acid (0.4 mol/L, 4.0 eq), HBTU (0.4 mol/L in DMF, 4.0 eq), HOBt (0.4 mol/L in DMF, 4.0 eq) and DIPEA (0.6 mol/L in DMF, 6.0 eq) were added to the reaction vessel and shaken for 1 h. After draining the coupling solution, the resin was consecutively washed with DMF, DCM (2 x) and DMF (4 x).

(2c) *N*-terminal acetylation (1.28 mmol scale) was achieved through reaction with acetic anhydride and DIPEA. Acetic anhydride (0.5 mol/L in NMP, 10.0 eq), HOBt (0.02 mol/L in NMP, 3.5 eq) and DIPEA (0.1 mol/L in NMP, 3.0 eq) were added to the reaction vessel and agitated for 10 min. After draining the reaction vessel, the previous step was repeated for 20 min. The resin was consecutively washed with DMF (2 x), DCM (2 x) and DMF (3 x).

(2d) Cleavage from the resin was performed through shaking the resin in TFE/DCM (1:4, 5 mL) for 45 min and precipitation in cold diethyl ether.

(3) Peptide-polymer coupling and deprotection

(3a) Telechelic H_2_N-PEO(3000)-NH_2_ (250.0 mg, 83 μmol, 1.0 eq) and the trityl-protected peptide Ac-FHFHFXG-OH (286.1 mg, 200 μmol, 2.4 eq) were dissolved in 5 mL DMF. PyBOP (112.8 mg, 217 μmol, 2.6 eq), HOBt (29.3 mg, 217 μmol, 2.6 eq) and DIPEA (58.1 μL, 333 μmol, 4.0 eq) were added. After stirring for 5 min at r.t., additional DIPEA (58.1 μL, 333 mol, 4.0 eq) was added. After 2 h of continued stirring, additional PyBOP (112.8 mg, 217 μmol, 2.6 eq) was added to the reaction mixture and stirring was continued over night at r.t. The volatiles were removed under reduced pressure and the residue dissolved in DCM. The reaction mixture was added dropwise to 40 mL of cold diethyl ether to precipitate the product.

(3b) After centrifugation and vacuum drying, TFA-mediated cleavage of the sidechain-protecting groups was performed: Cleavage solution (5 mL, TFA/TIPS/H_2_O 95:2.5:2.5) was added and shaken at r.t. for 45 min. After co-distilling with toluene three times, the residue was dissolved in DCM and precipitated in 40 mL of cold diethyl ether. Centrifugation and lyophilization gave the final product.

***Part II: Rheometric Characterizations:***

Hydrogels were formed on an *HR30 Discovery* Rheometer using a Peltier Plate and either an 8 mm flat plate geometry (*TA Instruments*) or a 20 mm cone and plate with 1° angle (*TA Instruments*). Gels were prepared by weighing out 1 wt % of the PPC and dissolving in acidic PBS (pH 2.4). Prior to loading gels on the rheometer, the PPC solution was brought up to a pH of 7 using 0.1 M NaOH. For experiments using the 8 mm flat geometry, 8 μL of solution were pipetted onto the Peltier plate and the geometry was lowered until the hydrogel filled the gap. For experiments using the 20 mm cone and plate geometry, 40 μL of sample was loaded onto the Peltier plate and the geometry gap was set to 24 µm per the geometry’s parameters.

***Part III: Cell Culture, Encapsulation, and Analyses***

Cell Expansion: CCL151 human pulmonary fibroblasts (ATCC, LL-24) were cultured using fibroblast basal medium (ATCC, PCS-201-030) and a serum-free fibroblast growth kit (ATCC, PCS-201-040) at a passage between 13-18.

Cell Encapsulation for 3D Culture: To encapsulate cells within the hydrogels for 3D cell culture, CCL151 cells cultured in tissue culture flasks were washed 1x with warm PBS, dissociated using 0.25% Trypsin-EDTA (*Thermo Fisher*, MT25053CI) and manually counted using a hemocytometer to prepare a solution of cells at 5x10^6^ cells per mL. Cells were pelleted via centrifugation (250 x g for 5 minutes). The gel solution was prepared by mixing 1 wt % PPC in acidic PBS (pH 2.4) followed by subsequently increasing the pH of the gel solution to 7 using 0.25 M NaOH. The cell pellet was resuspended in the gel solution, gel samples were pipetted into an 8 well chamber slide (*Thermo Fisher*, 155409) and incubated at 37 °C and 5% CO_2_ for at least 30 min. After 30 min, fresh serum-free complete fibroblast medium was pipette into the wells and cultures were left to incubate.

Live/Dead Assay: Viability of the cells in 3D culture at days 1, 3, and 5 was assessed using a live/dead cytotoxicity assay (*Thermo Fisher*, L3224). Briefly, samples with encapsulated cells were washed 1 x with sterile, warm PBS for 5 minutes and then incubated with calcein AM and ethidium homodimer per manufacturer’s instructions for 12 min at ambient temperature conditions. Hydrogels were washed 3 x with sterile warm PBS for 5 min each and imaged using an LSM 800 microscope (*Zeiss*) with 150 µm z-stacks such that at least 25 slices per sample were collected. All conditions were run in triplicate. Images were analyzed using Volocity (*PerkinElmer*) and the percentage of live cells was calculated using:

$$\frac{Total Live Cells (Green Cells)}{Total Cells (Green \& Red Cells)}*100\%=\% Live Cells$$

The total % of viable cells reported is an average of n=3 samples and standard deviation statistics were used to report error range.

Overnight Confocal Imaging of Reporter Cell Line: To assess 3D distribution of cells, a CCL151 lentiviral reporter cell line previously designed by the April Kloxin Lab was used.^1^ Briefly, cells were engineered to constitutively express a red fluorescent protein (DsRed) at all times, and conditionally express a green fluorescent protein (ZsGreen) upon upregulation of alpha smooth muscle actin, a well-known marker for myofibroblast activation. After 1 day in 3D culture, encapsulated reporter cells were imaged using an LSM 800 (*Zeiss*) confocal microscope with incubation at 37 °C and 5% CO_2_ for 17 hrs overnight. Confocol z-stacks (150 µm thick with at least 25 slices) were collected every 30 minutes for 17 hours and then compiled into an accelerated video to showcase encapsulation and motility in 3D culture.

***Part IV: Thermoresponsive cell retrieval proof of concept***

Primary Cell Encapsulation for 3D Culture and Cell Harvesting: To encapsulate cells within the hydrogels for 3D cell culture, CCL151 cells were cultured and passaged from cell culture flasks as prior mentioned. Hydrogel precursor solution was prepared by mixing 1 wt % PPC in acidic (pH 2.4) PBS followed by subsequently raising the precursor solution to pH 7 using 0.1 M NaOH. The cell pellet was then resuspended in the precursor solution and pipetted into individual wells of an 8-well chamber slide (Thermo Fisher, 155409) and incubated at 37 °C and 5% CO2 for hydrogel formation.

Owing to the thermoresponsive nature of the PPC hydrogels we hypothesized that fibroblasts could be recovered from these hydrogels without the use of any cell culture reagents. To accomplish this, 20 μL hydrogels with encapsulated cells were allowed to incubate and form for 2 hr. At harvesting, samples were removed from the incubator, and a small volume (~ 100 μL) of ice-cold PBS was added to the well, resulting in immediate dissolution of the hydrogel. Gentle pipette mixing was applied to ensure thorough dissolution of the hydrogel. Samples were then transferred to microcentrifuge tubes and pelleted at 2400 x rpm for 5 minutes in a high-speed microcentrifuge (Stellar Scientific, HS10050). The remaining liquid was then decanted from the cell pellet, and cells were resuspended in fresh serum-free culture media. For counting, a Trypan Blue exclusion assay was performed, and samples were counted and imaged using a Countess cell counter.

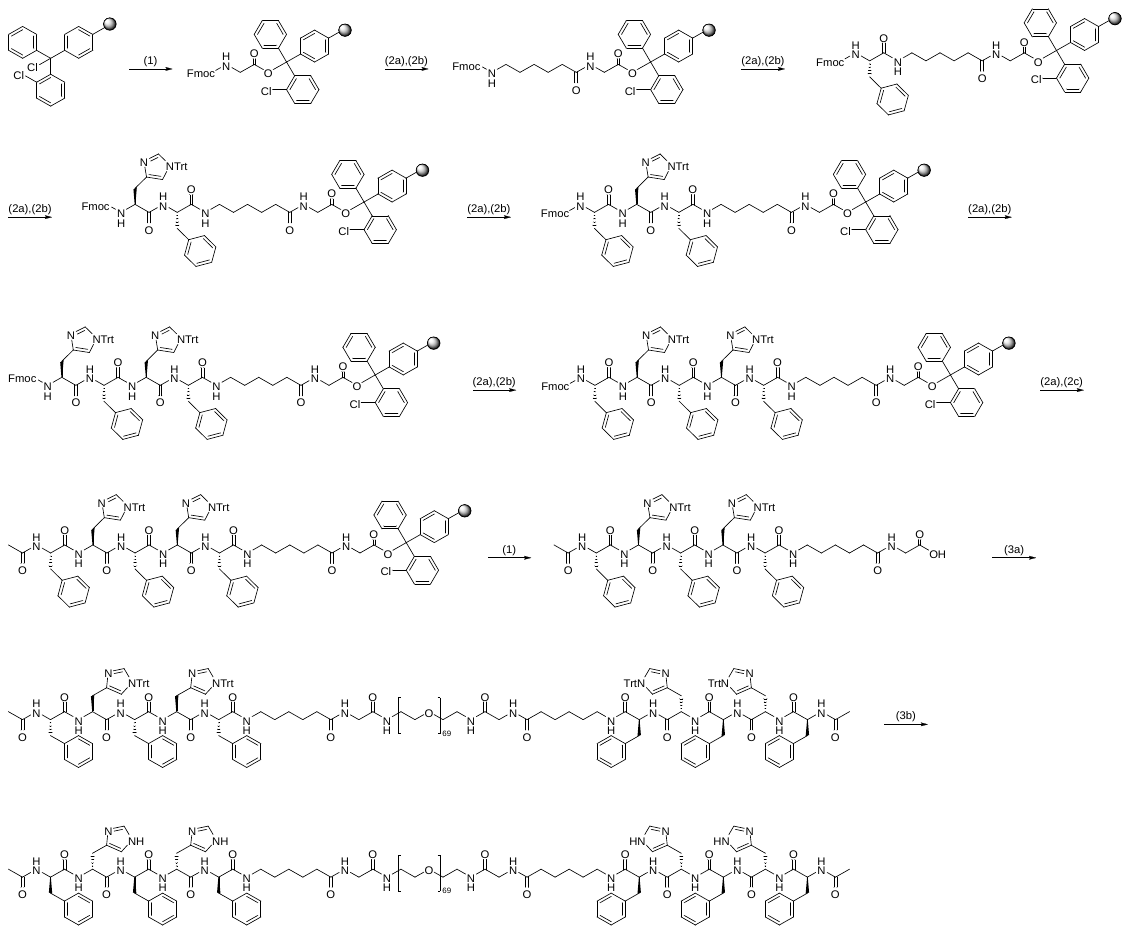

**Figure S1. Detailed schematic of polymer-peptide conjugate synthesis.

Yield:** 353.8 mg; 70.8 µmol; 85%; colorless solid.

**Molecular formula:** Calculated for n = 69: C_238_H_398_N_24_O_85_.

**Molecular weight:** Calculated for n = 69: 4955.83 g/mol.

**MALDI-ToF (Dit+KTFA, pos., linear mode) (*m/z*):** Calculated for n = 69 [C_238_H_398_N_24_O_85_K]^+^ = 4994.7937 g/mol, found: 4994.3420 g/mol

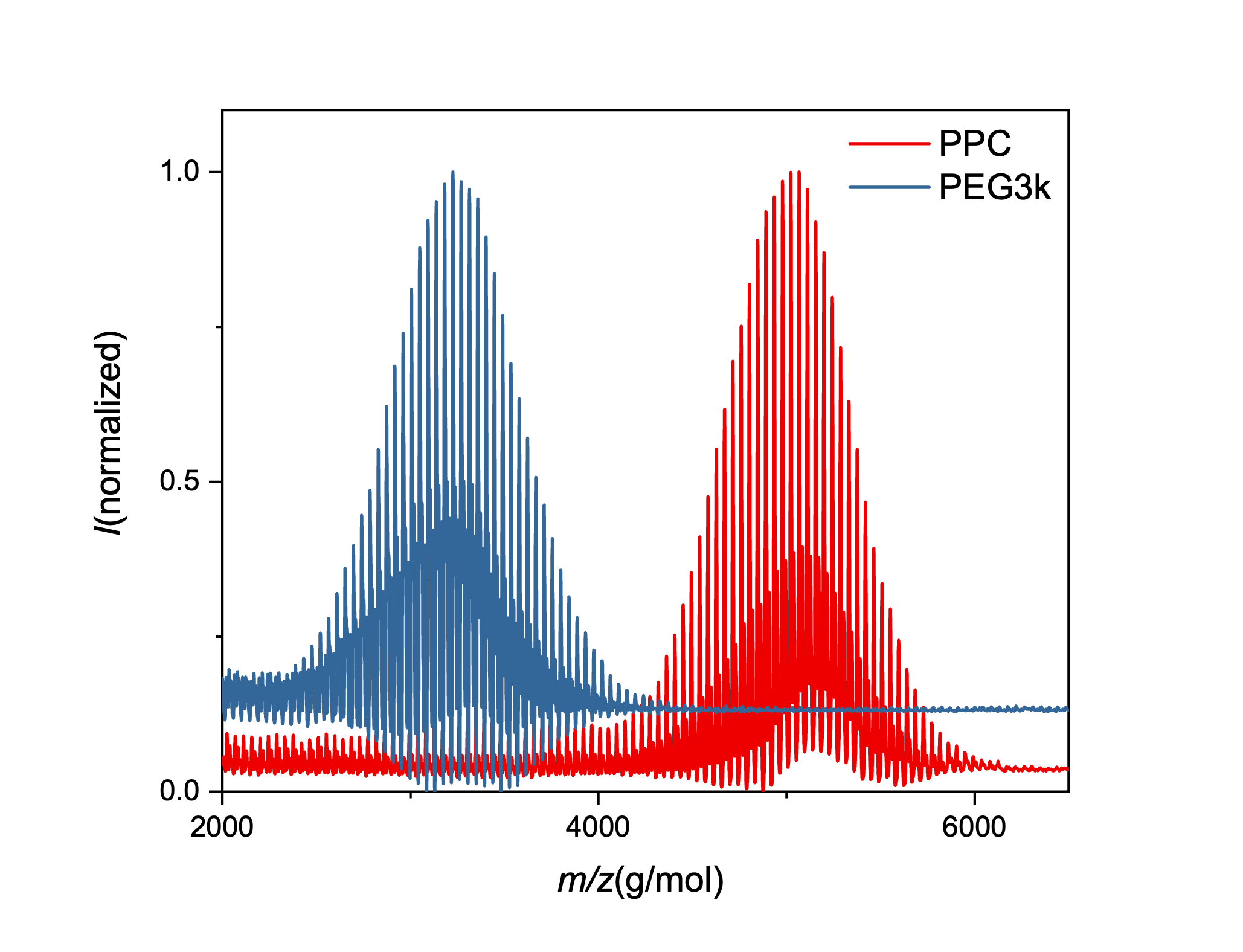

**Figure S2. MALDI-ToF-MS chromatogram of the PPC & PEG starting material**

**MALDI-ToF (Dit+KTFA, pos., linear mode) (*m/z*):** Calculated for n = 69 [C_238_H_398_N_24_O_85_K]^+^ = 4994.7937 g/mol, found: 4994.3420 g/mol

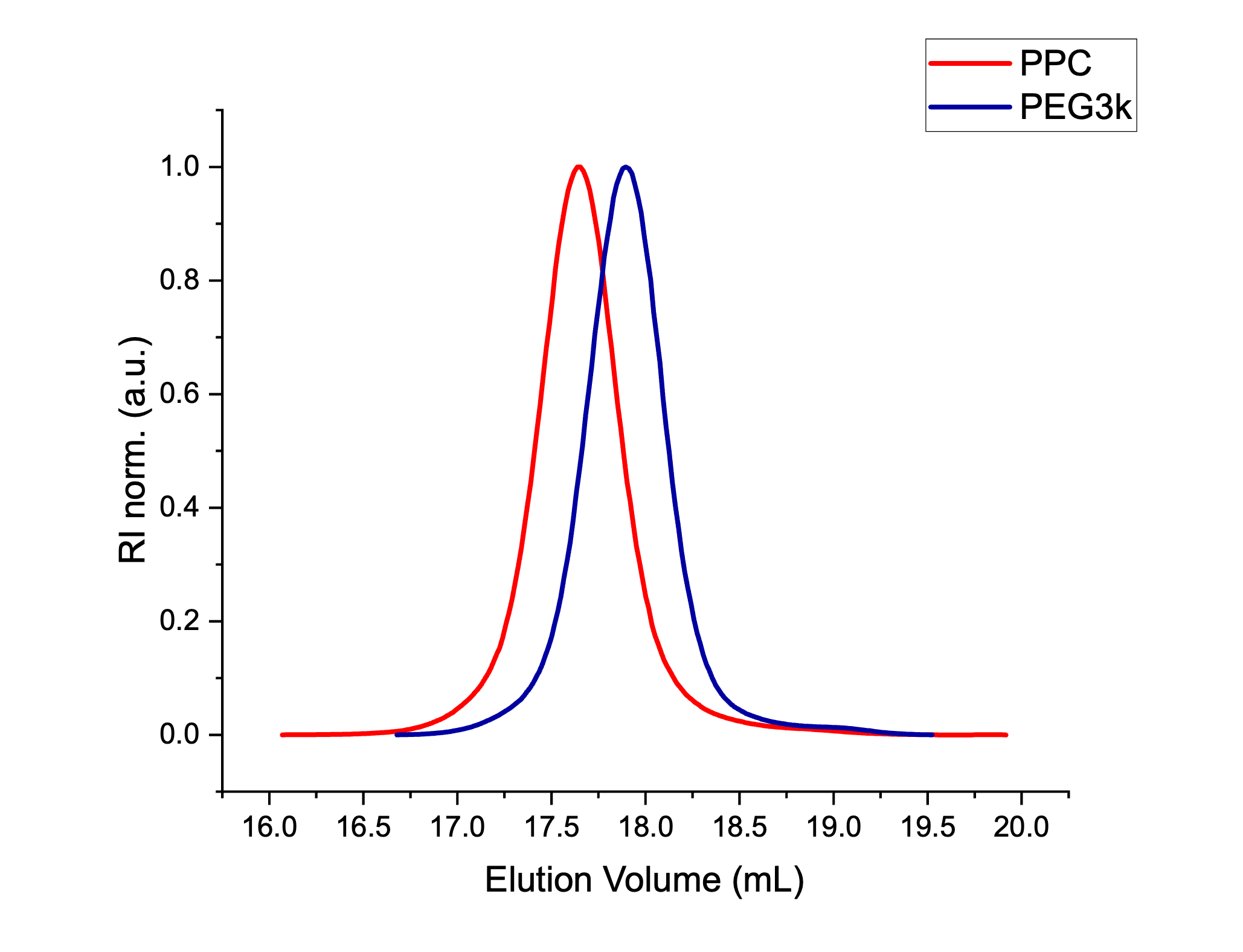

**Figure S3. MALDI-ToF-MS chromatogram of PPC**

Recorded in linear mode. Samples were prepared at concentrations of 1.0 mg/mL. Dithranol (Dit) was used as matrix with addition of potassium trifluoroacetate (KTFA).

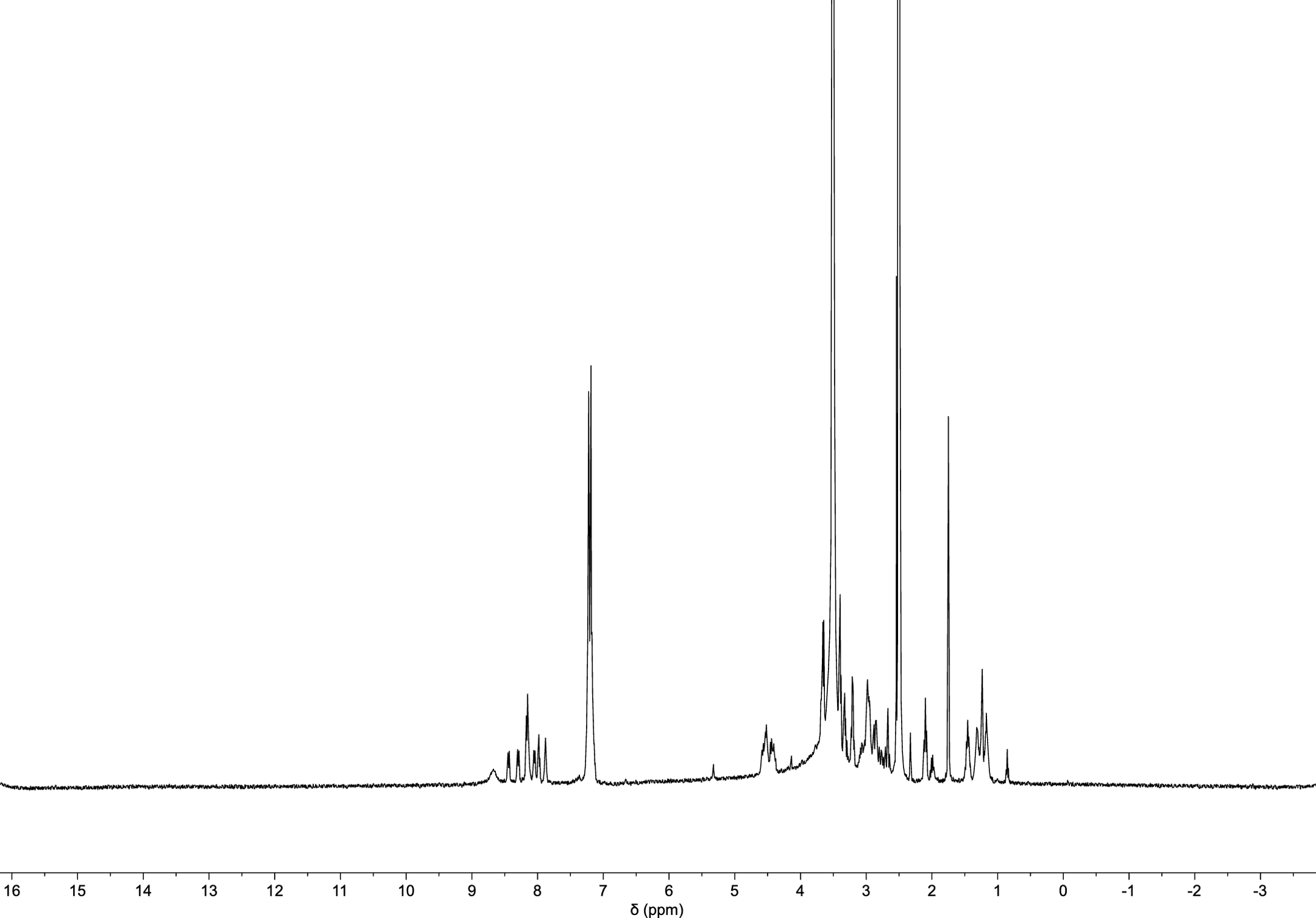

**Figure S4. ^1^H-NMR of PPC

^1^H-NMR (400 MHz, DMSO-*d_6_*):** δ [ppm] = 8.44 (d, 7.5 Hz, 3H, α-N**H**), 8.29 (d, 7.9 Hz, 2H, α-N**H**), 8.16 (d, 7.4 Hz, 8H, α-N**H**), 7.98 (t, 5.5 Hz, 2H, α-N**H**), 7.88 (t, 5.1 Hz, 2H, α-N**H**), 7.28-7.09 (m, 38H, C**H**^His^ / C**H**^Phe^), 4.49 (m, 18H, α-C**H**), 3.66 (m, 4H, C**H**_2_^Gly^), 3.50 (m, 276H, C**H**_2_^PEO^), 3.21 (q, 5.7 Hz, 10H, C**H**_2_^His^), 2.98 (s, 13H, C**H**_2_^Phe^), 2.72-2.61 (m, 8H, C**H**_2_^Ahx^) 2.10 (t, 7.4 Hz, 6H, C**H**_2_^Ahx^), 1.75 (s, 6H, C**H**_3_^NHAc^), 1.46 (t, 6.8 Hz, 6**H**, C**H**_2_^Ahx^), 1.31 (d, 7.5 Hz, 6**H**, C**H**_2_^Ahx^), 1.24 (s, 6**H**, C**H**_2_^Ahx^), 1.17 (m, 6**H**, C**H**_2_^Ahx^).

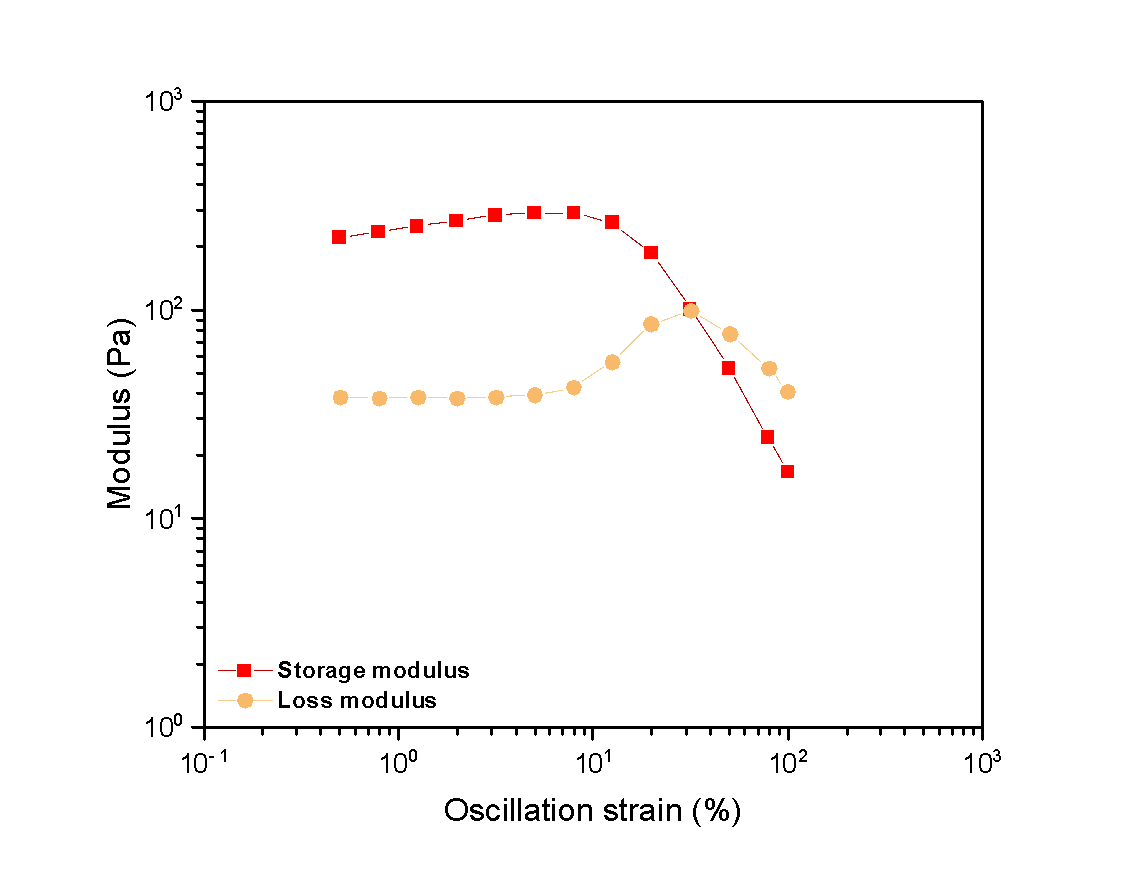

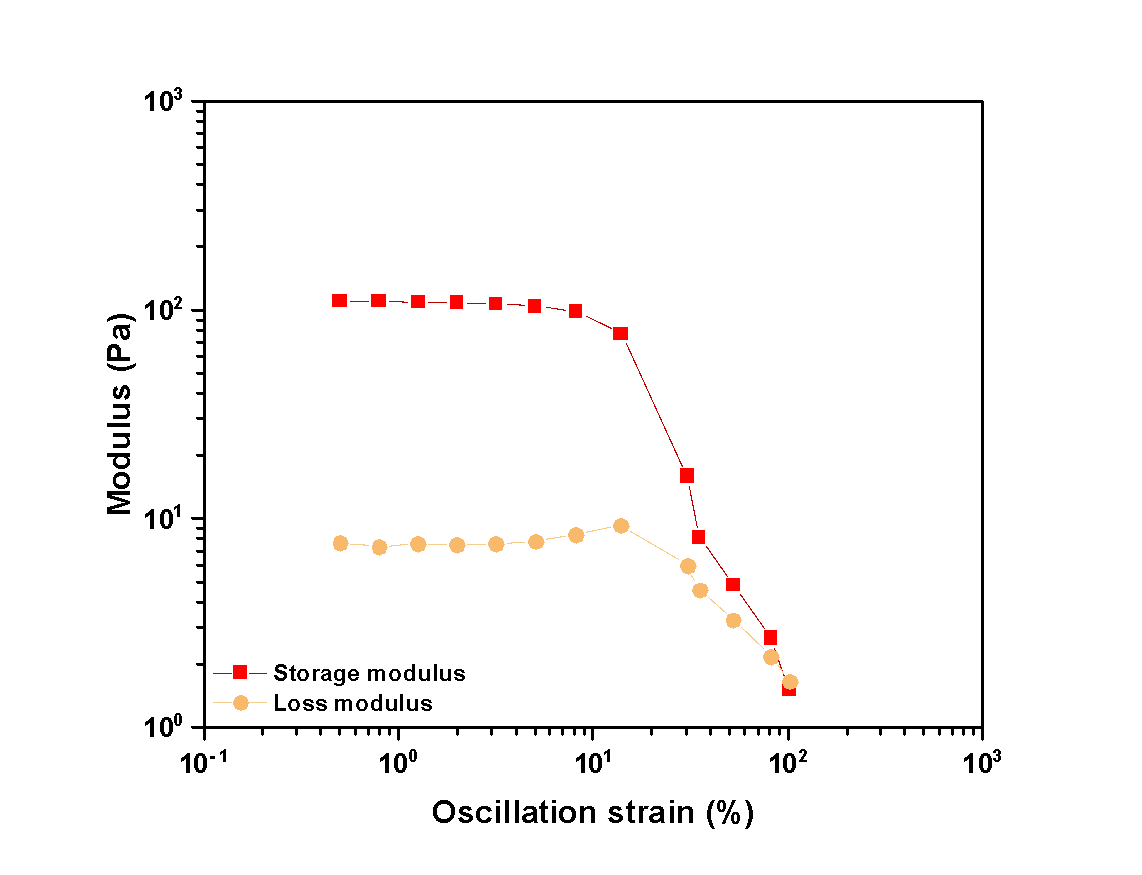

**Figure S5. Additional replicates of amplitude sweeps** to assess viscoelastic, shear thinning material properties.

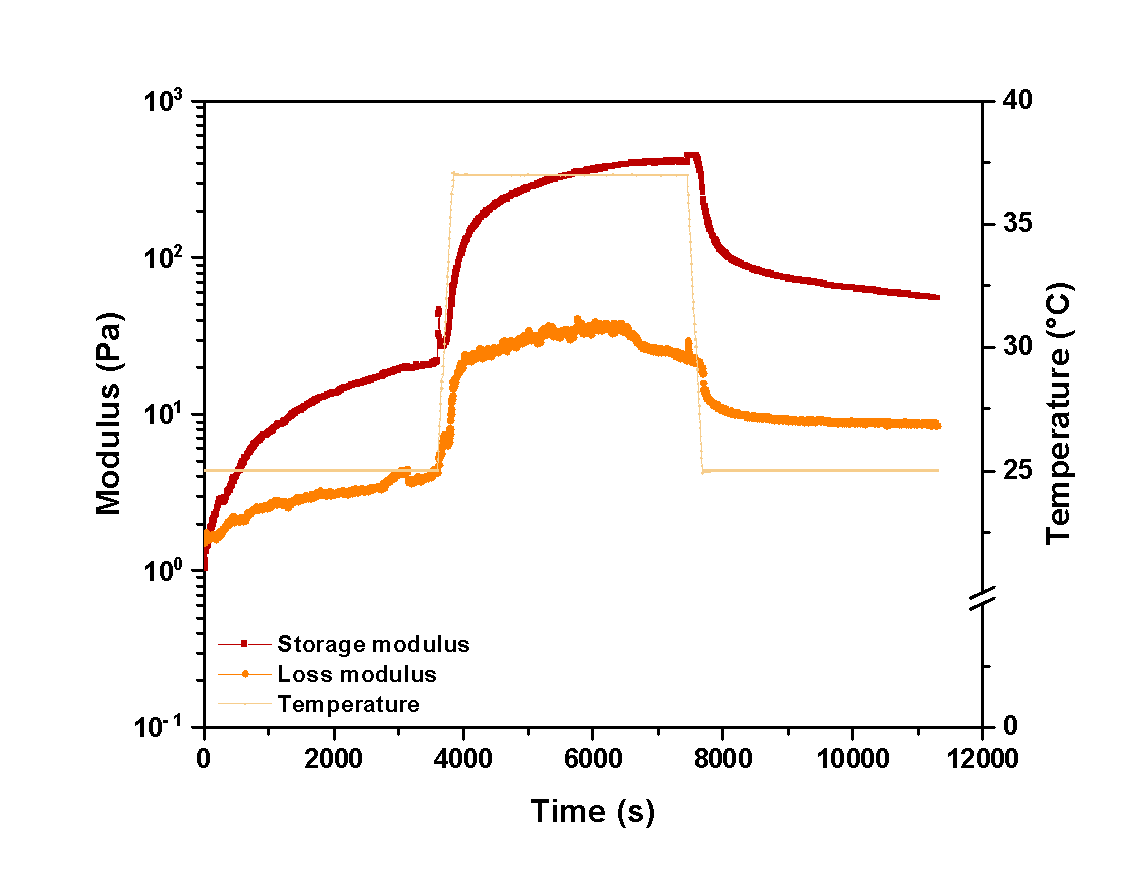

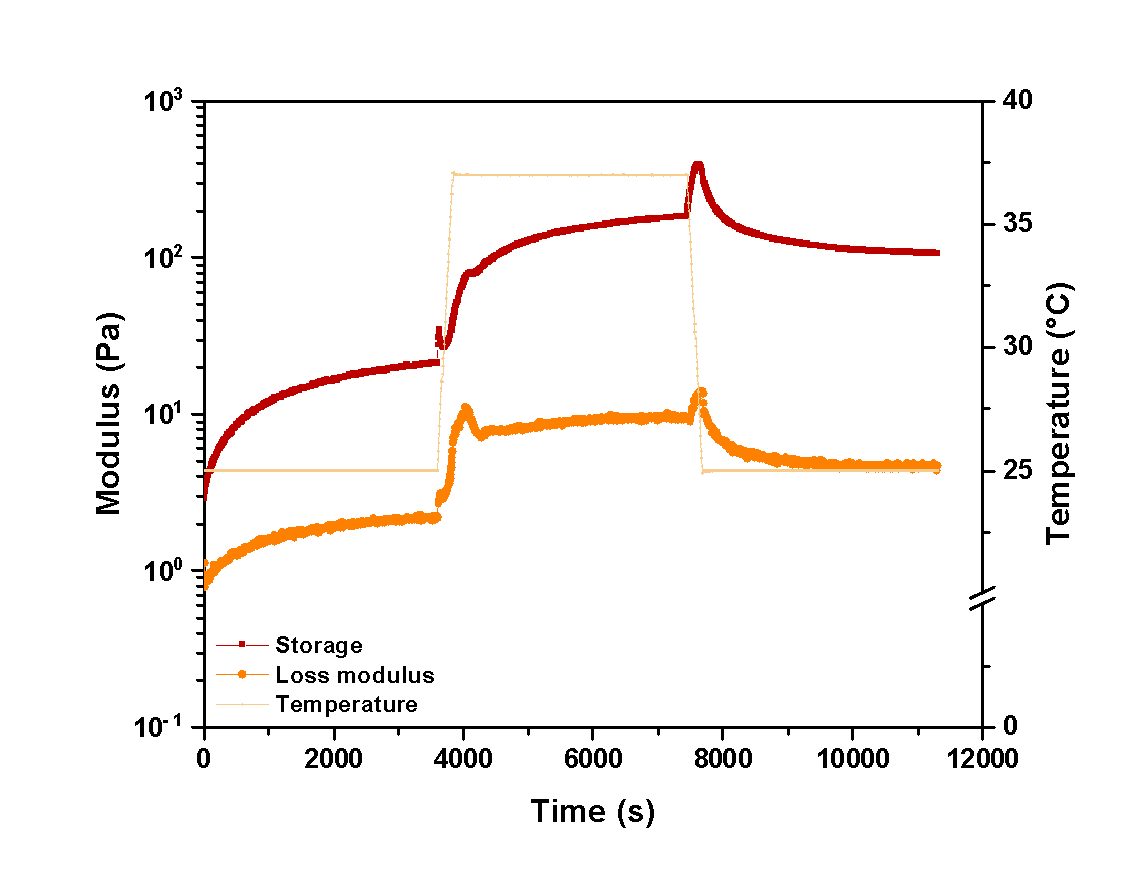

**Figure S6. Additional replicates of temperature ramp up studies** to assess material response to changes in environmental temperature.

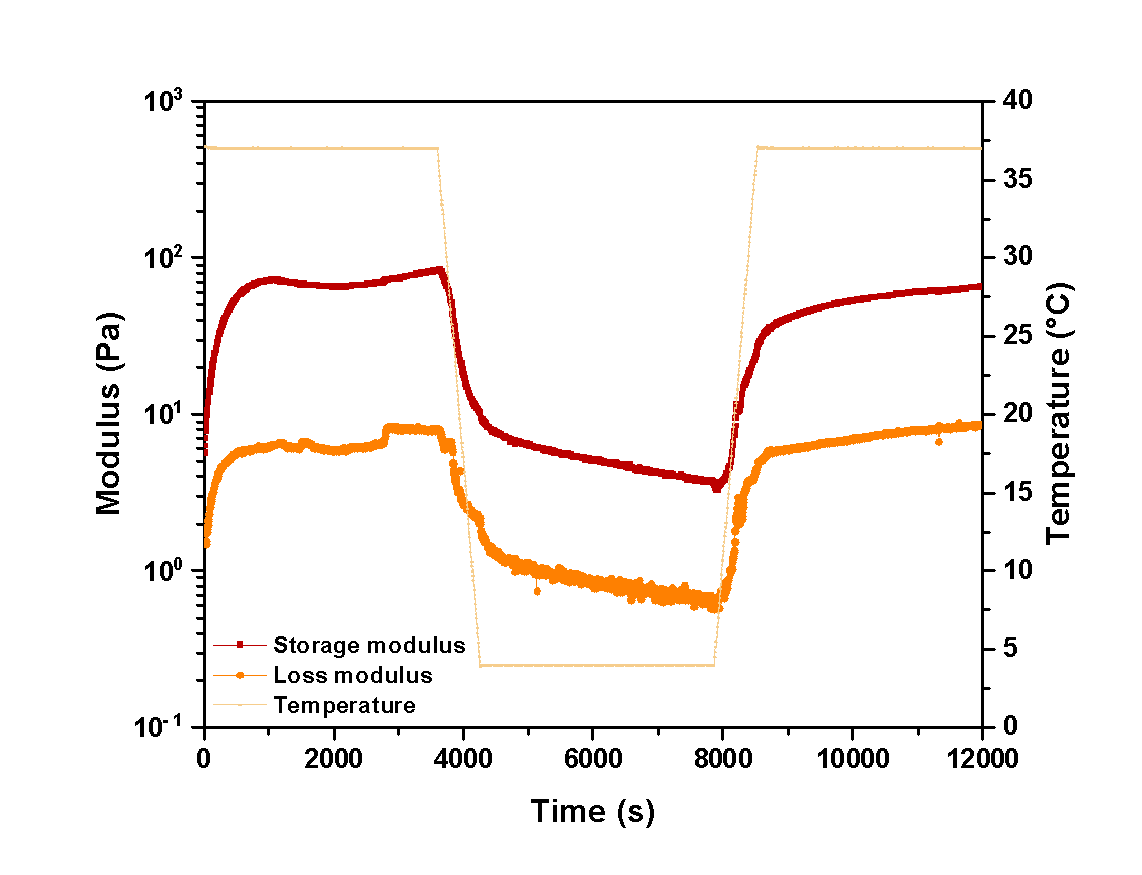

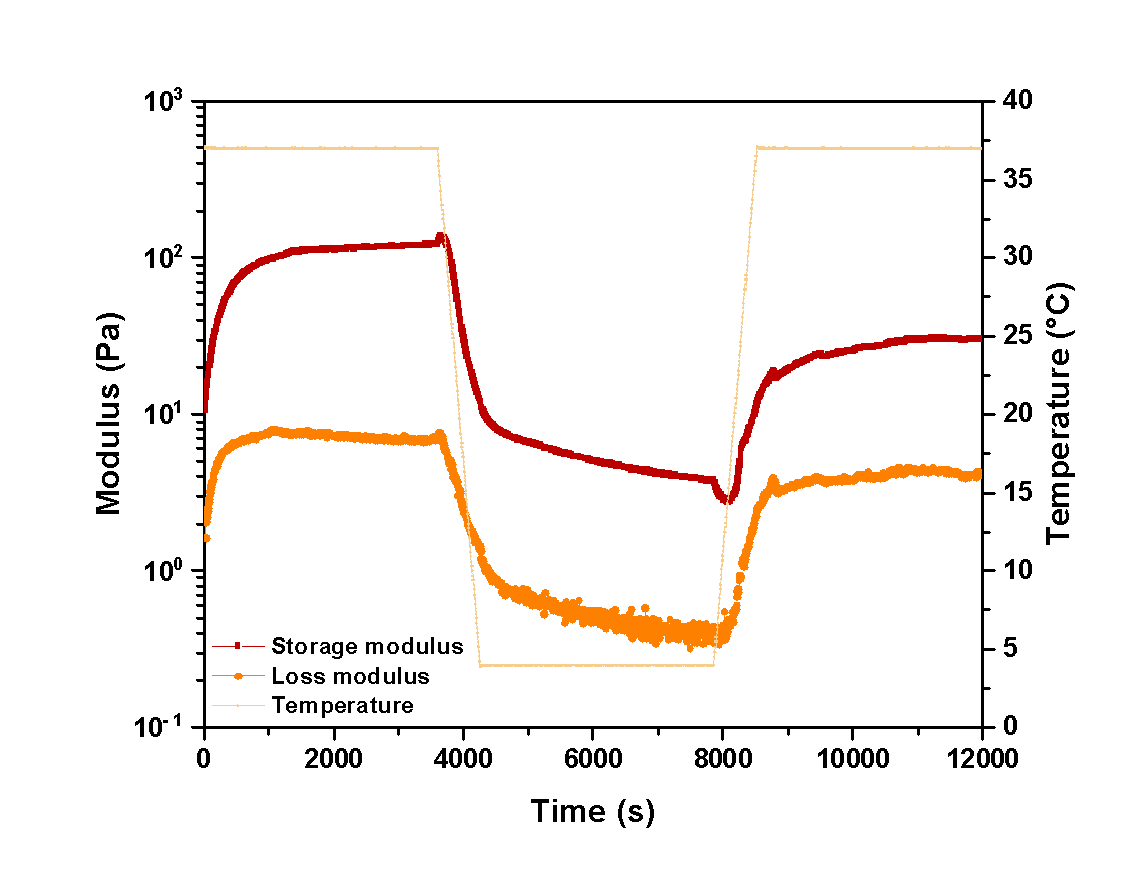

**Figure S7. Additional replicates of temperature ramp down studies** to assess material response to changes in environmental temperature.

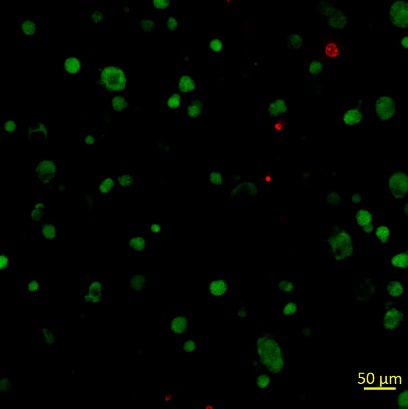

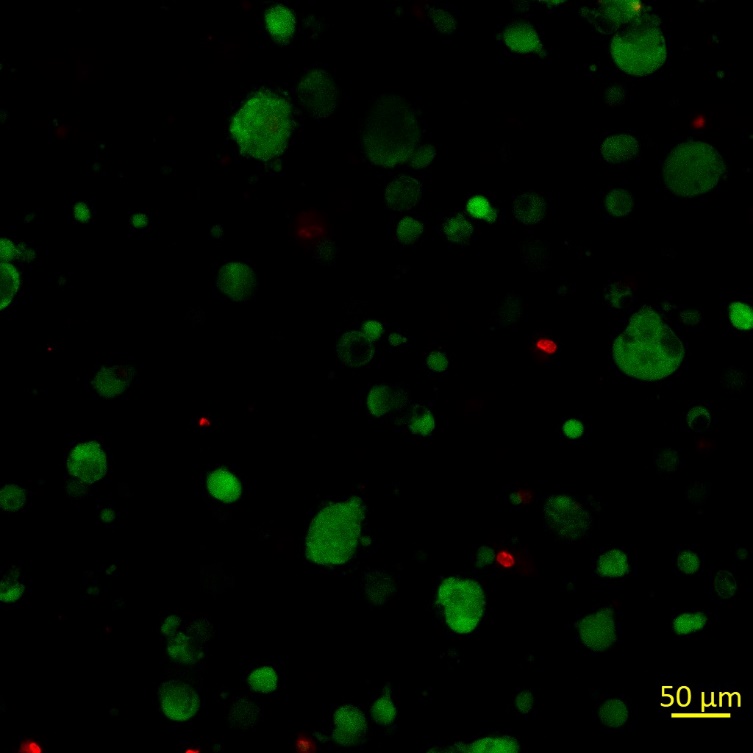

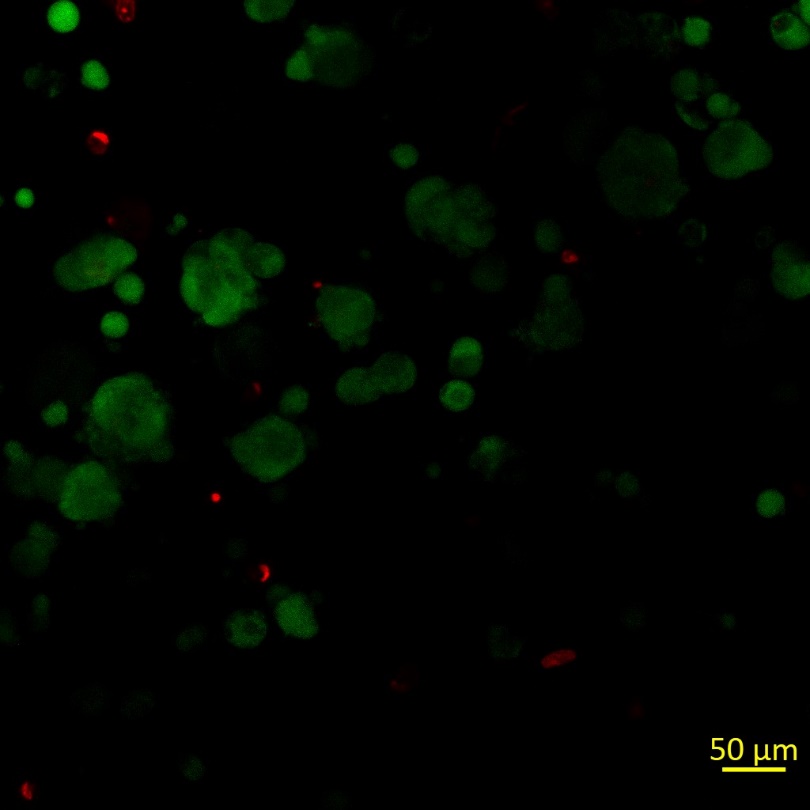

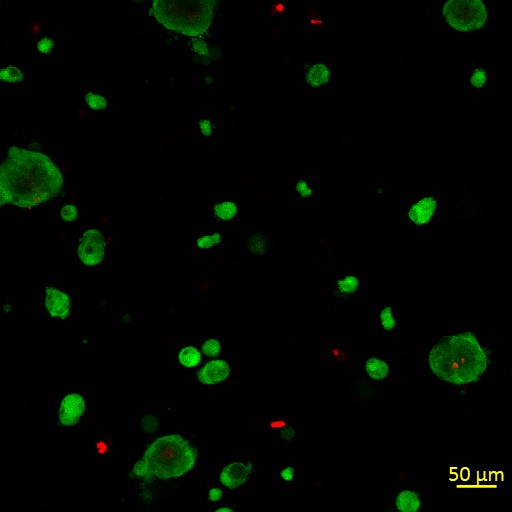

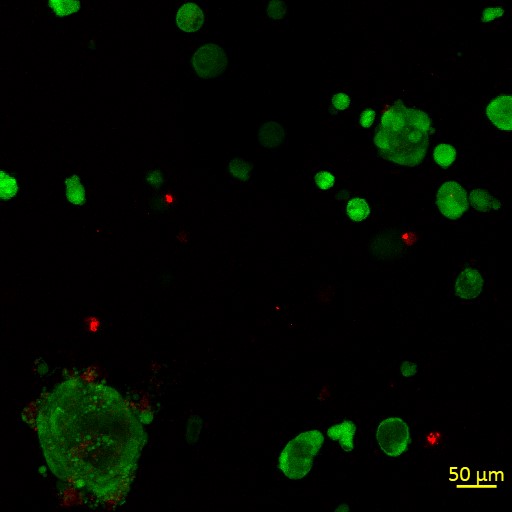

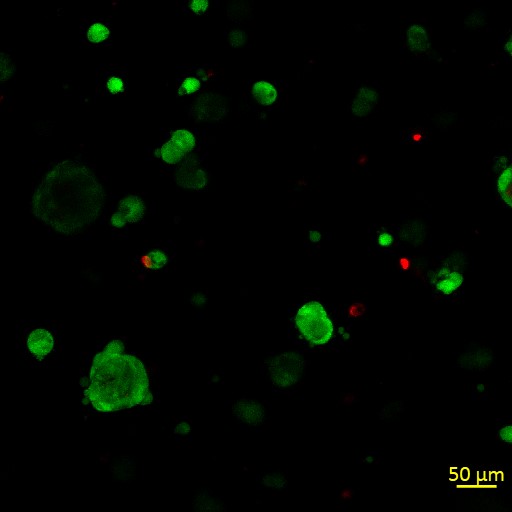

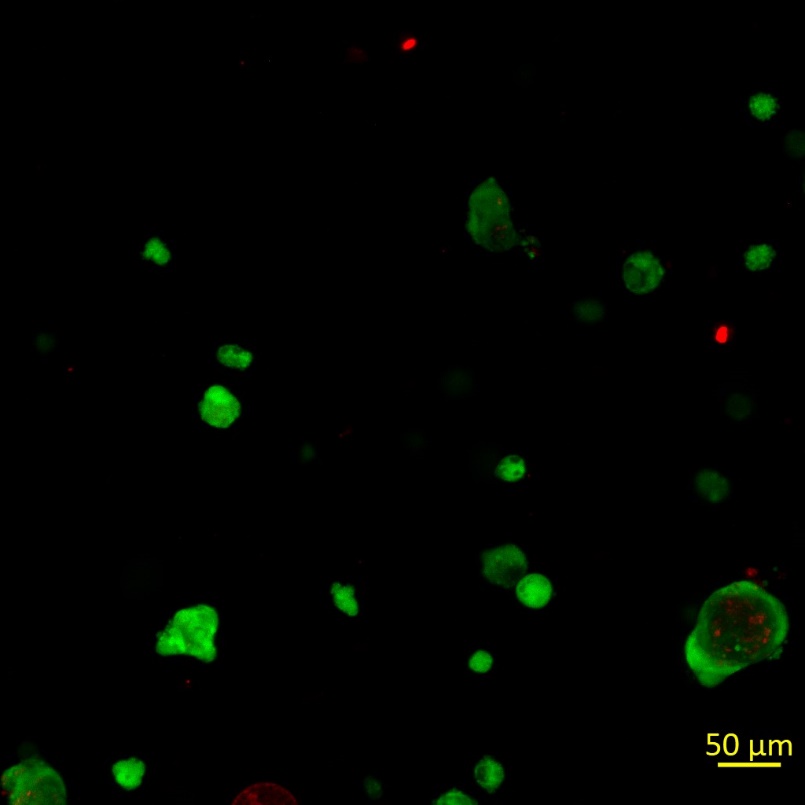

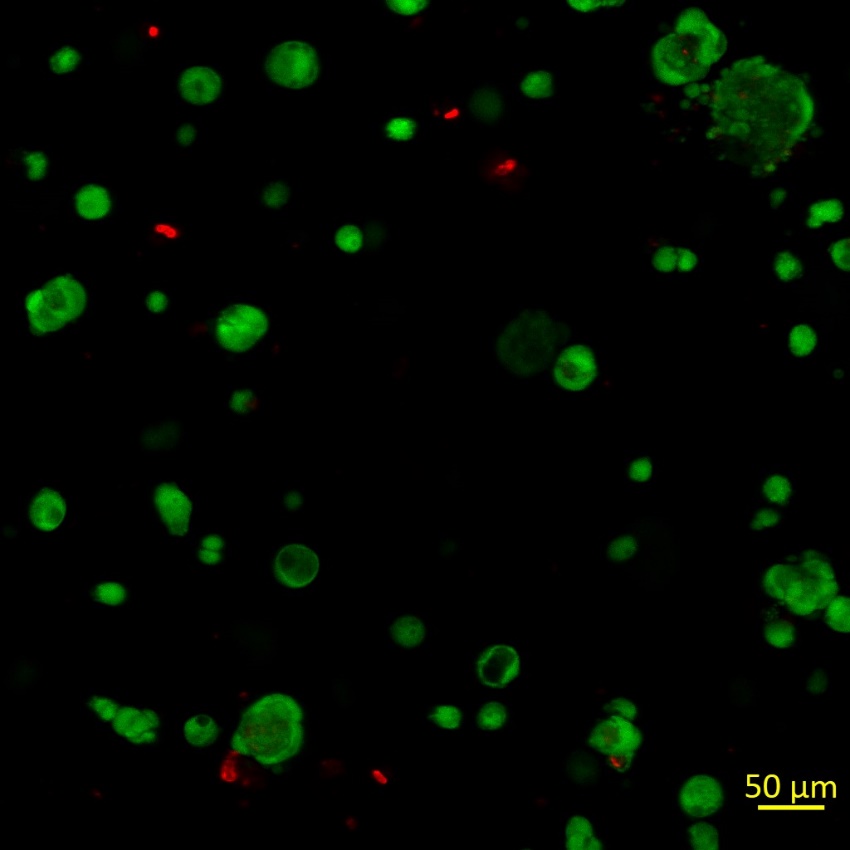

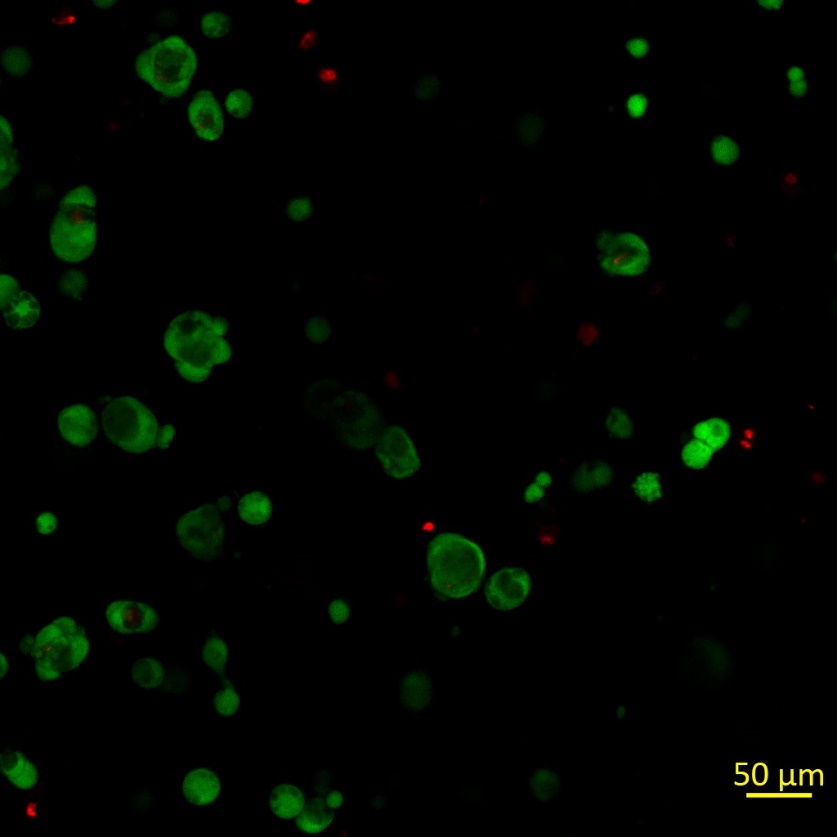

**Day 1**

**Day 3**

**Day 5**

**Replicate 1**

**Replicate 2**

**Replicate 3**

**Figure S8. All replicates of images used in cellular viability analysis.** Z-projections of confocal Z-stacks for assessment of cellular viability encapsulated within the hydrogels over a 5-day timespan shown.

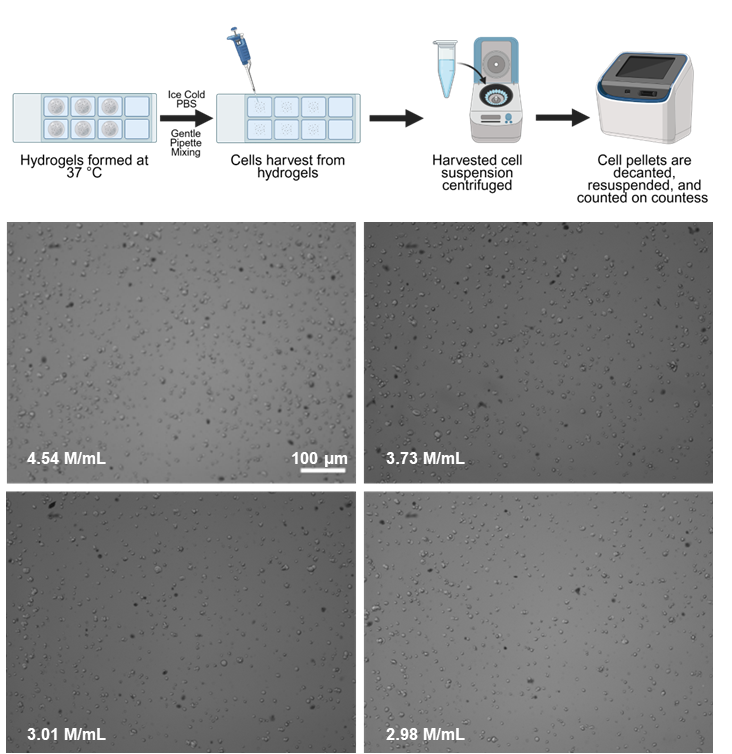

**E**

**D**

**C**

**B**

**A**

**Figure S9**. **Cell Harvesting.** A pilot study was performed for proof-of-concept for u thermoresponsive to harvest cells from the 3D cultures. (A) First, cells were suspended within the precursor solution (1 wt% PPC in PBS) and the cell suspension pipetted into individual wells of an 8-well chamber slide with hydrogel formation at 37 C for 2 hr. For cell harvesting from the 3D cultures, ice-cold PBS was then applied to the hydrogels of interest for their dissolution, and the resulting cell suspension within the wells was gently pipette mixed to further facilitate dissolution of any remaining hydrogel. Four hydrogel samples were pooled together to ensure a visible pellet was formed and a reliable cell count could be achieved. The harvested cell suspension was then centrifuged to pellet the cells (here, 2400 rpm was used on a Gusto High-Speed Mini centrifuge). The liquid over the cell pellet then was decanted, and the pelleted cells were resuspended in fresh serum-free growth media. At this stage, cells can be further processed or analyzed; here, we quantified cell number and viability using a Trypan Blue exclusion assay and a Countess cell counter. (B-E) Example Countess analysis of cell retrieval from 3D culture replicates. Based on this proof-of-concept, processing steps may be further optimized for other cell types or applications of interest.

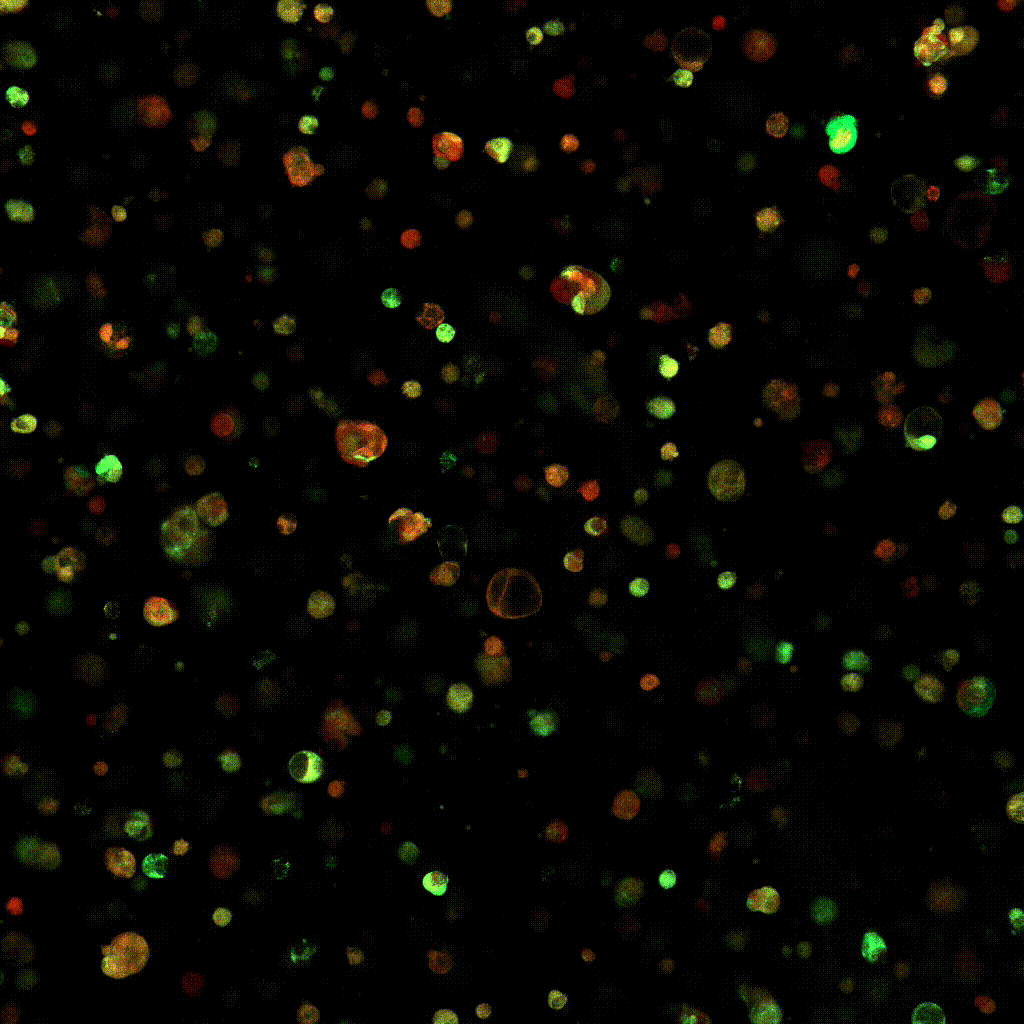

**Figure S10. Video of images collected of reporter line cells f**or 17 hours overnight 1 day after encapsulation within a hydrogel.
